# Single cell analysis of endometriosis reveals a coordinated transcriptional program driving immunotolerance and angiogenesis across eutopic and ectopic tissues

**DOI:** 10.1101/2021.07.28.453839

**Authors:** Yuliana Tan, William F. Flynn, Santhosh Sivajothi, Diane Luo, Suleyman B. Bozal, Anthony A. Luciano, Paul Robson, Danielle E. Luciano, Elise T. Courtois

## Abstract

Endometriosis is characterized by growth of endometrial-like tissue outside of the uterus affecting many women in their reproductive age, causing years of pelvic pain and potential infertility. Its pathophysiology remains largely unknown, limiting diagnosis and treatment. We characterized peritoneal and ovarian lesions at single-cell transcriptome resolution and compared to matched eutopic endometrium, control endometrium, and organoids derived from these tissues, generating data on over 100,000 cells across 12 individuals. We spatially localized many of the cell types using imaging mass cytometry. We identify a perivascular mural cell unique to the peritoneal lesions with dual roles in angiogenesis promotion and immune cell trafficking. We define an immunotolerant peritoneal niche, fundamental differences in eutopic endometrium and between lesions microenvironments, and a novel progenitor-like epithelial cell subpopulation. Altogether, this study provides a holistic view of the endometriosis microenvironment representing the first comprehensive cell atlas of the disease, essential information for advancing therapeutics and diagnostics.

## Introduction

Endometriosis is an inflammatory gynecologic condition that affects 6 to 10% of women in their reproductive age worldwide^1, 2^, with symptoms including but not limited to pelvic pain and infertility. It is characterized by the presence of endometrium-like tissue outside of the uterine cavity (termed lesions), commonly found within the peritoneal cavity, as superficial peritoneal or ovarian lesions. Endometriosis pathogenesis was proposed almost a century ago, and yet the exact etiology and molecular drivers of the disease remain largely unknown. Limited non-invasive diagnosis tools impede early detection, resulting in a gap of up to seven years from onset of symptoms to definitive diagnosis, which relies on invasive surgical biopsies of lesions. Current treatment of endometriosis remains similarly challenging, relying on hormonal therapy often in conjunction with invasive surgery for lesion removal. Oral contraceptives aim to reduce symptoms but do not necessarily promote lesion clearance. Even post-excision, lesions often recur, and repeated surgery is frequent^3^.

The struggle to diagnose and treat endometriosis is due in part to the poor understanding of the pathophysiology of, and heterogeneity within, endometriosis. It has been established that the immune system is implicated in endometriosis^4^, although its precise role remains poorly understood. The tissue microenvironment, including but not limited to immune cells, has been highlighted as a critical factor for normal development of the endometrium and disease progression in endometriosis^4–7^. Advancement in single-cell RNA sequencing (scRNA-seq) and organoid culture systems enables an understanding of the dynamic interactions within the microenvironment components of the endometrium and the cellular complexity and heterogeneity in endometriosis. Recent studies have demonstrated the power of such cutting-edge technologies to understand the human endometrium and how this dynamically changes through the menstrual cycle and pregnancy^5–7^.

In this study, we profiled and analyzed the transcriptome of endometrium and endometriotic lesions at the single-cell level using scRNA-seq and hyperplexed antibody imaging. To understand the changes in endometrium and endometriotic lesions during treatment, our cohort focuses on patients who are experiencing active disease symptoms while undergoing oral contraceptive treatment. Through profiling eutopic endometrium, two types of endometriosis lesions (peritoneal and ovarian), and patient-derived organoids, we uncover both distinct cellular changes in endometriosis endometrium as well as specific subsets of immunomodulatory macrophages, immunotolerant dendritic cells (DCs), and vascular changes unique to endometriosis. Our data highlight a novel endometriosis-specific pericyte population, as well as an unreported progenitor-like epithelial cell population which may be critical for deeper understanding of this disease.

## Results

### Single cell transcriptomics and imaging mass cytometry (IMC) reveal the cellular heterogeneity of eutopic endometrium and ectopic lesions and highlight major changes in cell composition

Single cell RNA-seq (scRNA-seq) was performed on biopsies from a total of 12 individuals. Control eutopic endometrium (Ctrl) represented samples from non-endometriosis patients. Eutopic endometrium (EuE), ectopic peritoneal lesions (EcP) and the adjacent regions to these (EcPA), and ectopic ovarian lesions (EcO) were collected from ASRM Stage III-IV endometriosis patients (Fig. 1a; Supplementary Fig. 1a, Supplementary Table 1). The EcPA was included in order to study the environment where lesions establish and evolve. In total, 90,414 single cell transcriptomes were generated, with an overall median of 9,950 unique transcripts and 2,994 genes per cell (Fig. 1b). Cells were assigned to one of five overarching cell types: epithelial, stromal, endothelial, lymphocyte, and myeloid (Fig. 1c,d). Iterative clustering within each of these broad cell classifications resulted in the identification of 58 subpopulations (Fig. 1d, Supplementary Table 2), highlighting the cellular complexity of both the endometrium and ectopic lesions. In order to understand what, if any, biases were introduced by our tissue dissociation process, we compared the bulk transcriptomes from undissociated tissue to pseudo-bulk single cell transcriptomes (Supplementary Fig. 1a). As expected, differential expression analysis for each tissue type indicates that adipocytes, neuronal projections, and muscle cells are not well represented in our single cell dataset (Supplementary Fig. 1b, Supplementary Table 3). Nevertheless, transcriptome similarities between bulk and single cell data indicate that our single cell dataset reflects much of the overall original tissue composition and its cellular complexity.

**Fig. 1.**
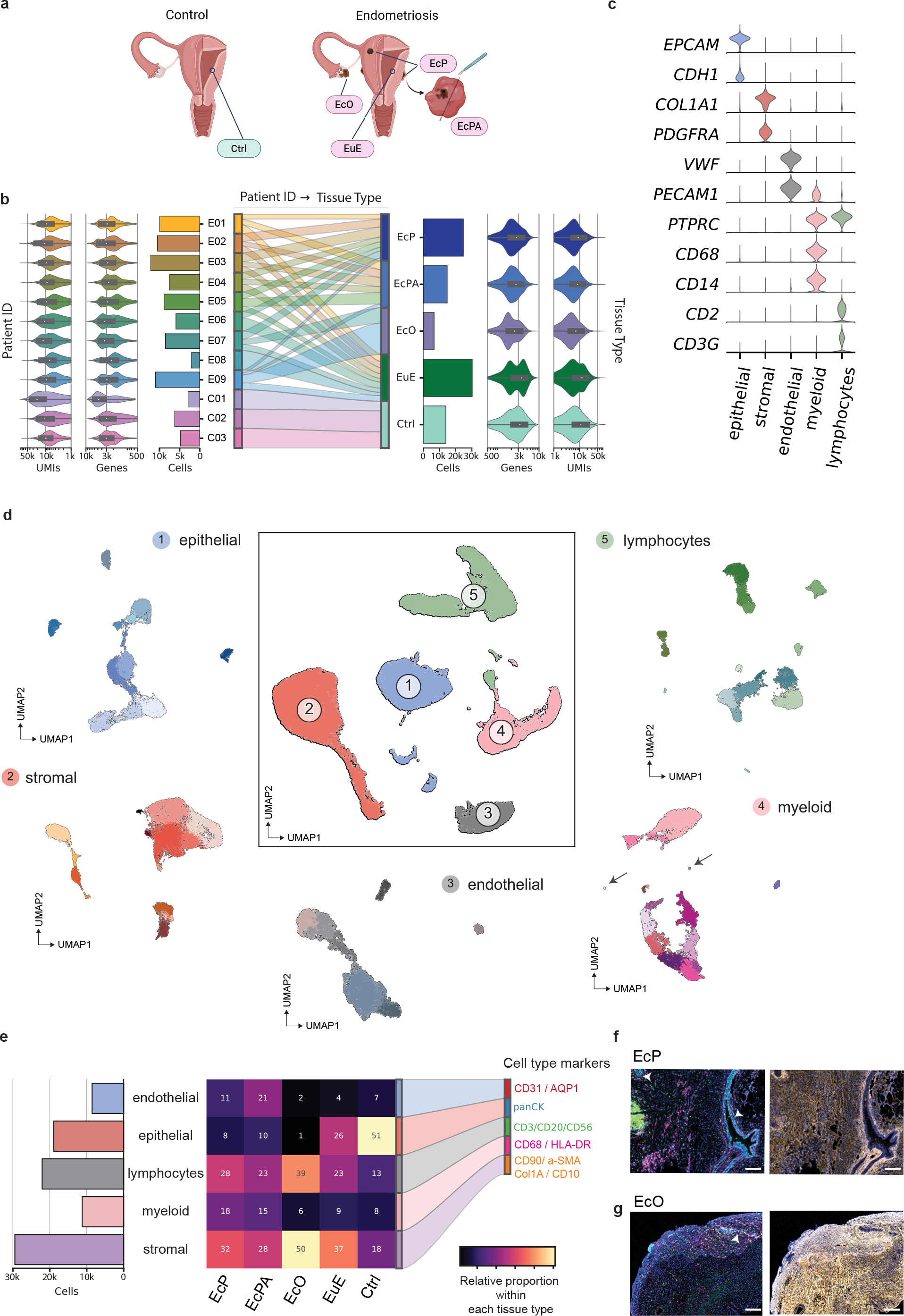
scRNA-seq from control and endometriosis patient. **a**, Schematic representation of collected tissue biopsies. Control (Ctrl) specimens were obtained from eutopic endometrium of women without endometriosis. Eutopic endometrium (EuE), ectopic peritoneal endometriosis (EcP), ectopic peritoneal adjacent lesion (EcPA), and ectopic ovarian endometriosis (EcO) were obtained from women with endometriosis. **b**, Diagram showing scRNA-seq metrics per patient (left) and tissue type (right) after QC. These metrics indicate unique molecular identifier (UMIs) and total genes per cells across patients and tissue types. The cord diagram (center) indicates how each patient (E: endometriosis, C: control) contributes to each tissue types. **c**, Violin plot representing marker gene expression for each major cell type identified in the scRNA-seq dataset. **d**, UMAP plot showing the 90,414 single-cells from control and endometriosis tissues. Five major cell types are identified (center UMAP plot) and subsequently subclustered into 58 subpopulations (radial UMAP plots). Each subpopulation was identified using marker genes curated form the literature. The presence of basophils and neutrophils (arrows) indicate that the cell recovery workflow was suited to capture delicate cell types, known to be easily lost during tissue dissociation. **e**, Diagram showing number of major cell types (bar plot) and the proportion for each tissue type (heatmap plot). Cell proportions are indicated within each square. Unique combinations of cell markers from each major cell cluster were used to design an IMC panel. Assigned colors represents each major cell type identified in EcP (**f**) and EcO (**g**). White arrows indicate endometriotic epithelial glands. Scale bar = 100 µm.

The heterogeneity among the profiled tissues is evident in the changes in cell type composition. First, the two endometriosis lesion types, EcP and EcO, display markedly different cellular proportions (Fig 1e). Second, EcP and EcPA are highly similar, particularly among epithelial cells, which suggests that endometriosis lesions may extend beyond the macroscopic core of the lesions and into the peritoneum. Third, the composition of eutopic endometrium in EuE differs dramatically relative to Ctrl, with much of the epithelial component replaced by lymphocytes and stroma in EuE. Consistent with this finding, we observed an increase in the expression of cell cycle-related genes and proliferation of endometrial fibroblasts in EuE (Supplementary Fig. 2a-c). On a per-patient basis, we find that EuE biopsies stratify into two groups distinguished by immune cell or fibroblast abundance, both distinct from Ctrl samples (Supplementary Fig. 2d). This indicates endometriotic endometrium has distinct transcriptomic signatures that differ from normal endometrium and this is created by changes in cell type composition. This composition change is a result of an increase in relative contributions of fibroblast or immune cells, and points towards the heterogeneity in eutopic endometrium across individual patients.

Through spatial cell analysis we aimed to determine how cellular organization mediates unique and specific cell signaling pathways that could explain the endometriosis associated changes observed in the single cell datasets. We designed an antibody panel to visualize each of the major cell types and subpopulations identified by scRNA-seq analysis and performed (IMC) on peritoneal and ovarian lesions (Supplementary Fig. 3). Through IMC we confirmed the scarcity of epithelial glands and predominant presence of stromal cells within the EcO when compared to EcP (Fig. 1f,g), validating our single-cell compositional analysis above.

### Active vascular remodeling and immune cell-trafficking prevail in endometriosis peritoneal lesions

Among the changes in cellular composition detected in endometriosis, we found endothelial cells (EC) to be markedly increased in peritoneal lesions, suggesting a possible role for angiogenesis. We identified components of the vascular system: four mural and seven endothelial cell subpopulations, according to careful analysis of previously described marker gene expression^5,6,8–10^(Fig. 2a-c). Mural cells, which include vascular smooth muscle cells (VSMC) and perivascular cells (Prv), are specialized cells that directly interact with ECs to provide support and promote blood vessel stabilization. Mural cells account for roughly 50% of the stromal cells in EcPA (Supplementary Fig. 4a), suggesting a highly vascularized microenvironment. An increased relative proportion of endothelial cells in EcPA also supports this (Fig.1e). Prv-STEAP4 and Prv-MYH11 were previously identified in the endometrium^6^, but we also observed a novel Prv-CCL19 subpopulation expressing both *STEAP4* and *MYH11*. This subpopulation accounts for the majority of pericytes in EuE, EcP and EcPA, is absent in EcO, and exhibits tissue specific gene expression patterns (Supplementary Fig. 4b-d). We find Prv-CCL19 with increased abundance and *CCL19* expression specifically in and around peritoneal lesions together with upregulation of known angiogenesis regulators such as Synuclein-y (*SNCG)*^11^ and angiopoietin genes (*ANGPT1*, *ANGPTL1* and *ANGPT2*)^12^. Similarly, Prv-CCL19 upregulate expression of ligands implicated in T-cell recruitment^13^ (*CCL21* and *FGF7*) (Fig. 2d-e). Interestingly, *SUSD2*, a marker for endometrial mesenchymal stem cells identified in endometriosis^14^, is specifically co-expressed in EcP and EcPA *CCL19*^+^ Prv cells (Supplementary Fig. 4d). Together, these data show the presence of an endometriosis specific perivascular subpopulation, likely promoting angiogenesis and immune chemotaxis in peritoneal lesions (Fig. 2f).

**Fig. 2.**
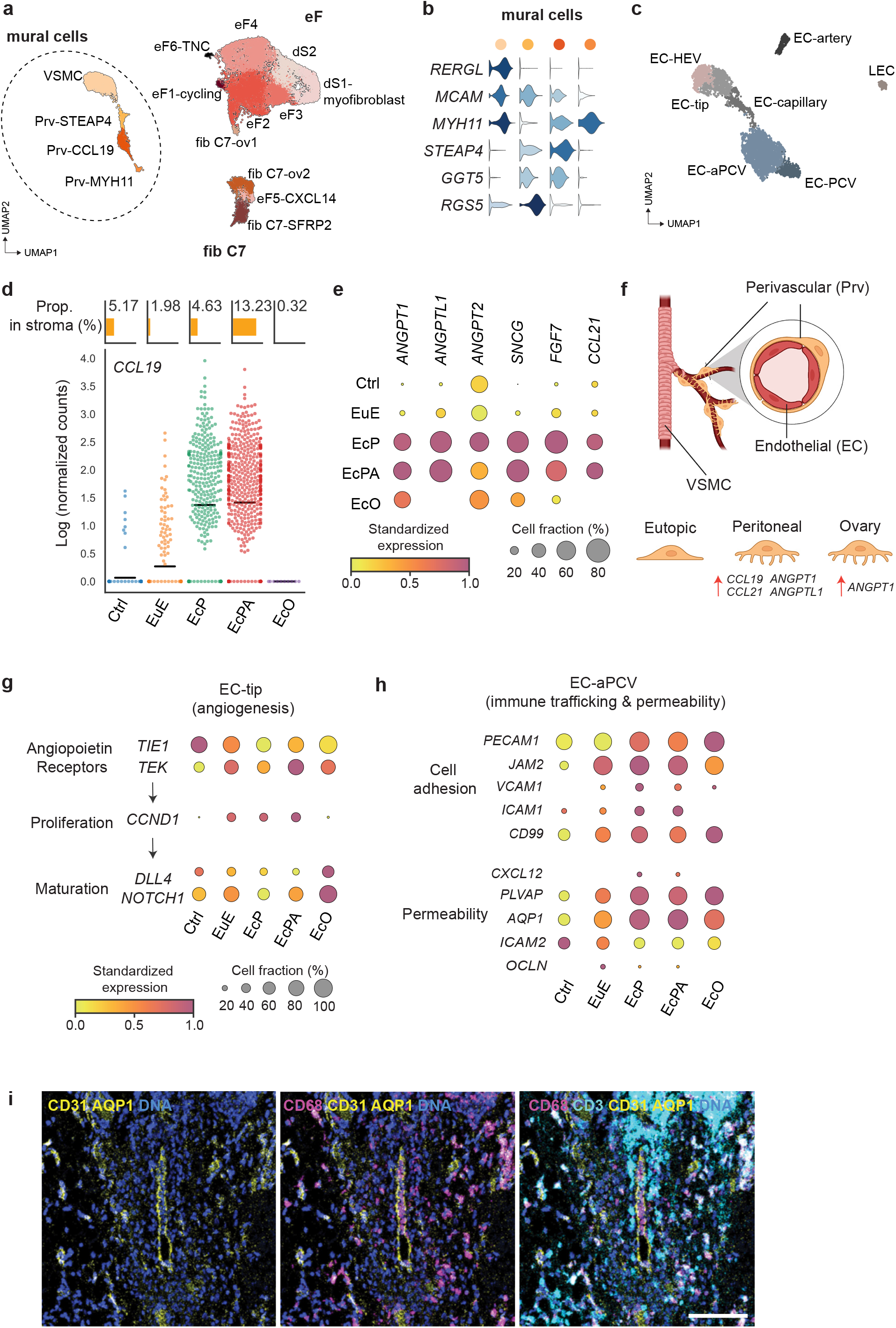
Role of Stromal cell diversity in angiogenesis and immune trafficking in endometriosis lesions. **a**, UMAP plot of stromal cells representing the 15 identified subpopulations and classified into three general cell subtypes: endometrial fibroblast (eF), fibroblast C7 (fib C7) and mural cell (n= 28,751 cells). **b**, Violin plot showing markers of mural cell subpopulations. **c**, UMAP plot of endothelial cells (EC), represented across 7 subclusters: lymphatic EC (LEC), high endothelial venule (EC-HEV), tip EC (EC-tip), capillary (EC-capillary), post-capillary vein (EC-PCV), activated PCV (EC-aPCV), and arterial (EC-artery). **d**, (top) Proportion of Prv-CCL19within stromal cells. A major increase of Prv-CCL19 is observed in EcPA. Bars represent the mean value. (bottom) The swarm plot shows *CCL19* expression in individual cells from each lesion. **e**, Dot plot showing significantly upregulated genes involved in angiogenesis and immune cell trafficking (edgeR, FDR < 0.05) in Prv-CCL19. **f**, Schematic of mural and EC localization. Larger arteries and veins are unsheathed by VSMC, while smaller vessels (e.g., capillaries) are unsheathed by perivascular cells. Lesions Prv cells increase expression of pro-angiogenic genes when compared to Ctrl. **G**, Dot plot showing significant representative DEGs involved in new vessel formation in tip EC (edgeR, FDR <0.05). **h**, Dot plot showing significant DEGs involved in cell adhesion and permeability in a-PCV (edgeR, FDR <0.05). **I**, Representative IMC image from a peritoneal lesion. CD3^+^ T-cells (cyan) and CD68^+^ myeloid cells (magenta) localize near blood EC vasculature marked by CD31 and AQP1 (yellow). Nuclei counterstained by DNA labeling (blue). Scale bar = 100µm.

To further elucidate the interactions between ECs and Prv-CCL19, we performed ligand-receptor analysis using a modified workflow based on CellPhoneDB (See Methods); our data indicate that EC-tip cells respond to the *ANGPT1* produced by pericytes, an interaction known to induce tube formation and branching^15^. In endometriosis, and specifically in EcPA, *TEK* expression is upregulated while expression of *TIE1*, the anti-angiogenic receptor for angiopoietins^12, 16^, is downregulated (Fig. 2g; Supplementary Table 4). Previous studies have shown that *TEK* pathway activation leads to EC proliferation and activation of a feedback loop mechanism through *DLL4-NOTCH1* signaling to induce tip EC maturation^17^. We found a significant increase in the expression of cell cycle genes and a decrease in *DLL4* expression in EuE and peritoneal lesion (EcP and EcPA) relative to control endometrium (Fig. 2g), suggesting higher proliferative capacity of tip ECs. On the other hand, Notch signaling is upregulated in EcO, suggesting the presence of more mature tip ECs in the ovarian microenvironment.

Immune cell trafficking involves extravasation of immune cells from the blood stream into the interstitial tissue, where ECs act as the barrier. Extravasation mainly occurs at the capillaries and post-capillary venous (PCV) level^18^. Activated PCV (EC-aPCV) and EC-PCV cells proportions, two subpopulations of PCVs^9, 10^, are remarkably increased in ectopic lesions and adjacent peritoneal tissue (Supplementary Fig. 5a). We found that genes which regulate immune cell attachment and monocyte trafficking^18^—*PECAM1*, *JAM2*, *VCAM1*, *ICAM1*, *CD99*—and genes associated with EC permeability^19^— *PLVAP, AQP1, CXCL12*—are upregulated in endometriosis EC-aPCV, while genes encoding tight junction proteins responsible for endothelial-to-endothelial cell contact^18^—*ICAM2 and OCLN––*are downregulated (Fig. 2h). We find, through IMC, abundant lymphocytes (CD3^+^) and myeloid (CD68^+^) cells both within and surrounding blood vessels (marked by CD31 and AQP1), indicating active immune trafficking at this site (Fig. 2i). Expression of *AQP1,* which also associates with increased angiogenesis and migration of endothelial cells^20^, is substantially increased in endometriosis EC-PCVs and EC-aPCVs (Supplementary Fig. 5b,c). These data indicate the presence of a leaky PCV vasculature in peritoneal lesions, allowing increased immune cell chemotaxis.

Thus, our data describe an endometriosis-specific perivascular subtype and emphasize on the dynamic orchestration of Prv-CCL19 and PCV endothelial subpopulations to promote angiogenesis and immune cell trafficking in peritoneal endometriosis lesions. This analysis also highlights some substantial differences between ovarian and peritoneal lesions microenvironments.

### Macrophage subtypes contribute to the neurogenic, angiogenic, and immunosuppressive microenvironment of ectopic peritoneal and ovarian lesions

Our scRNA-seq analysis uncovered 16 myeloid cell subpopulations (Fig 3a, Supplementary Fig. 6a) and 14 lymphocyte subpopulations (Fig. 1d). Myeloid cells, and in particular macrophages (Mϕs), have been characterized as central components of the endometriosis ecosystem, playing a key role in the establishment of endometriosis^2^. This, together with our observations indicating an increase in myeloid cell presence in peritoneal lesions (Fig. 1e) prompted us to investigate the complexity of the macrophage population across different tissues. We identified five Mϕ subpopulations, of which Mϕ1-LYVE1 and Mϕ3-APOE were previously identified by single cell analysis in other systems^8, 21, 22^. Tissue-resident macrophage subpopulations (Mϕ1-LYVE1, Mϕ2-peritoneal and Mϕ3-APOE) are distinguished by their expression of *FOLR2—*a gene associated with embryonic-derived tissue resident macrophages^19, 23^. Mϕ2-peritoneal subpopulation is exclusive to peritoneal tissue and express *ICAM2*, a known marker for peritoneal macrophage^24^. Mϕ4-infiltrated cells are present in all tissues and express *CLEC5A*, *CCR2*, and *VEGFA*, all markers for blood infiltrated macrophages^25, 26^ (Fig. 3b,c). In order to elucidate the putative relationships between these Mϕ populations, we performed RNA velocity trajectory analysis and hierarchical clustering on mean expression patterns of each population. Both suggest that Mϕ3-APOE are more similar to tissue resident Mϕ1-LYVE1 and Mϕ2-peritoneal subpopulations, as opposed to blood infiltrated Mϕ4 cells that appear more similar to monocytes (Fig. 3d, Supplementary Fig. 6b). Interestingly, Mϕ5-activated cells, characterized by activation markers, appear to arise from both infiltrated and tissue-resident macrophages (Fig. 3b-d). Together, these data illustrate the presence of distinct tissue-resident and blood infiltrated macrophage populations in endometrial tissue.

**Fig. 3.**
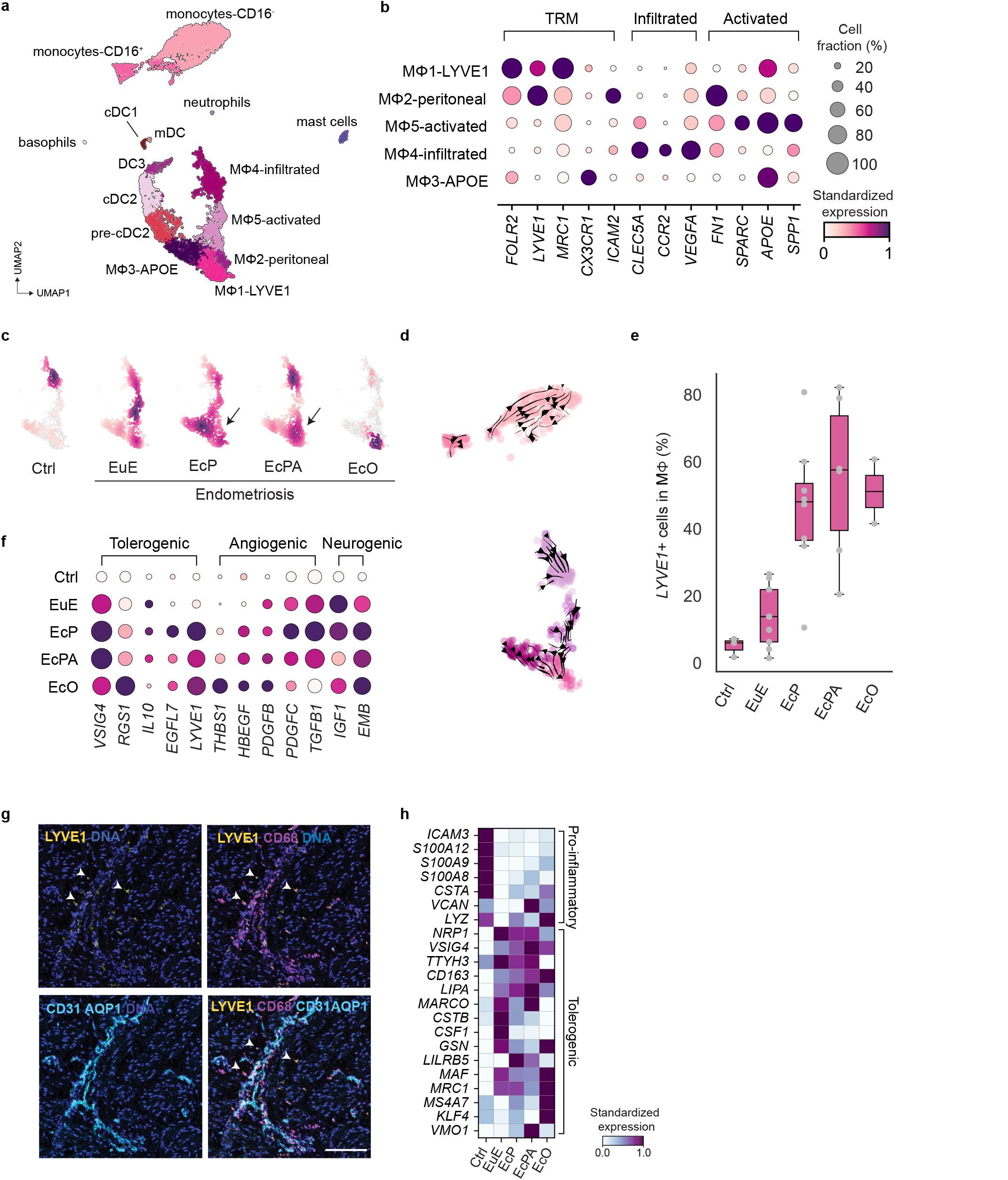
Macrophage heterogeneity in control and endometriosis. **a**, UMAP plot of myeloid cells, clustered into 15 different subtypes (n= 11,113 cells) (pDC are not included in this UMAP). **b**, Dot plot showing expressed genes in tissue-resident (TRM), blood-infiltrated, and activated macrophages across Mϕ identified subpopulations across all tissues. **c,** Density plot showing macrophage distribution for in each tissue type. **d,** UMAP plot showing RNA velocity streamlines for monocytes and macrophages in Ctrl. Streamlines represent the predicted transition path of cells across the subpopulations. **e,** Bar plot showing the proportion of *LYVE1-*expressing cells to all macrophages within each tissue type. Each dots represent percentage of LYVE1+ cells in a tissue biopsy. The box represents quartiles of the dataset with centerline corresponding to median and whiskers spans the min and max data point. **f,** Dot plot showing DEG involved in immunotolerance in Mϕ1-LYVE1 population. **g,** IMC image from FFPE tissue section of a peritoneal lesion. Images depict LYVE1^+^ macrophages (LYVE1, CD68) localization near endothelial cells (CD31, AQP1) (white arrows). Scale bar =100µm. **h,** Matrix plot showing expression of pro-inflammatory and pro-tolerogenic related genes in Mϕ4 subpopulation in Ctrl and endometriosis in.

Different macrophage subpopulations are enriched in endometriosis eutopic endometrium and ectopic tissues, respectively, and the relative proportion of macrophage subtypes dramatically altered between control and endometriosis. Mϕ1-LYVE1 and Mϕ5-activated abundance is substantially increased in EuE and EcP compared to Ctrl (Fig. 3c, Supplementary Fig. 6c). Most macrophages present in EcO belong to the Mϕ1-LYVE1 subtype, markedly distinct from the peritoneal lesion macrophage landscape. Across patients, *LYVE1^+^* macrophages are enriched in both eutopic and ectopic endometriosis tissues compared to Ctrl (Fig. 3e). Tissue resident *LYVE1^+^* macrophages have been previously associated with angiogenesis^19, 21^, arterial stiffness^27^ and anti-inflammatory phenotypes^23^. In agreement, we found that endometriosis Mϕ1-LYVE1 upregulate tolerogenic (*IL10*, *VSIG4*, *RGS1*, *EGFL7*) and angiogenesis-related genes (*THBS1*, *HBEGF*, *PDGFB*, *PDGFC, IGF1*) (Fig. 3f). Furthermore, inflammation and antigen presenting pathways of Mϕ1-LYVE1 are downregulated in endometriosis tissues (Supplementary Table 5) and Mϕ1-LYVE1 localization along the blood vessel*—*not within blood vessel*—*confirms the link between angiogenesis and this specific cell population (Fig. 3g). In addition, two genes (*IGF1*, *EMB*) previously shown to promote neurogenesis sprouting in endometriosis^28^ and neuromuscular junctions^29^, were among the top upregulated genes in Mϕ1-LYVE1 in endometriosis tissues, in both eutopic and ectopic, suggesting their implication in pain-related mechanism (Fig. 3f). The Mϕ4-infiltrated population, which is almost completely absent in EcO, presented with pro-tolerogenic features in endometrial tissue in stark contrast to its pro-inflammatory presentation in control endometrium (Fig. 3h). Altogether, we identified endometriosis-associated changes in multiple macrophage subpopulations, promoting tolerogenic, pro-angiogenic and pro-neurogenic microenvironment. Moreover, the altered macrophage landscape of lesions is also observed in endometriosis eutopic endometrium, affecting both tissue resident and blood infiltrated macrophages.

### Dendritic cells from the peritoneal lesion periphery adopt an immunomodulatory phenotype

Among the six populations of DCs we identified, we classified three *CD1C^+^* populations as pre-cDC2, cDC2, and DC3 according to previously reported marker genes^22, 30, 31^ (Fig 4a). We found the DC proportions vary greatly across patients (Supplementary Fig. 7a). However, *CD1C*^+^ DCs consistently accounted for the majority of the DCs in all tissues (Fig. 4b). Ovarian lesion DC populations differed substantially from peritoneal lesions, showing an increase in cDC1s and a dramatic reduction of cDC2s (*CD1A*^+^ DCs), again illustrating another difference between peritoneal and ovarian endometriosis lesion (Fig. 4b-d).

**Fig. 4.**
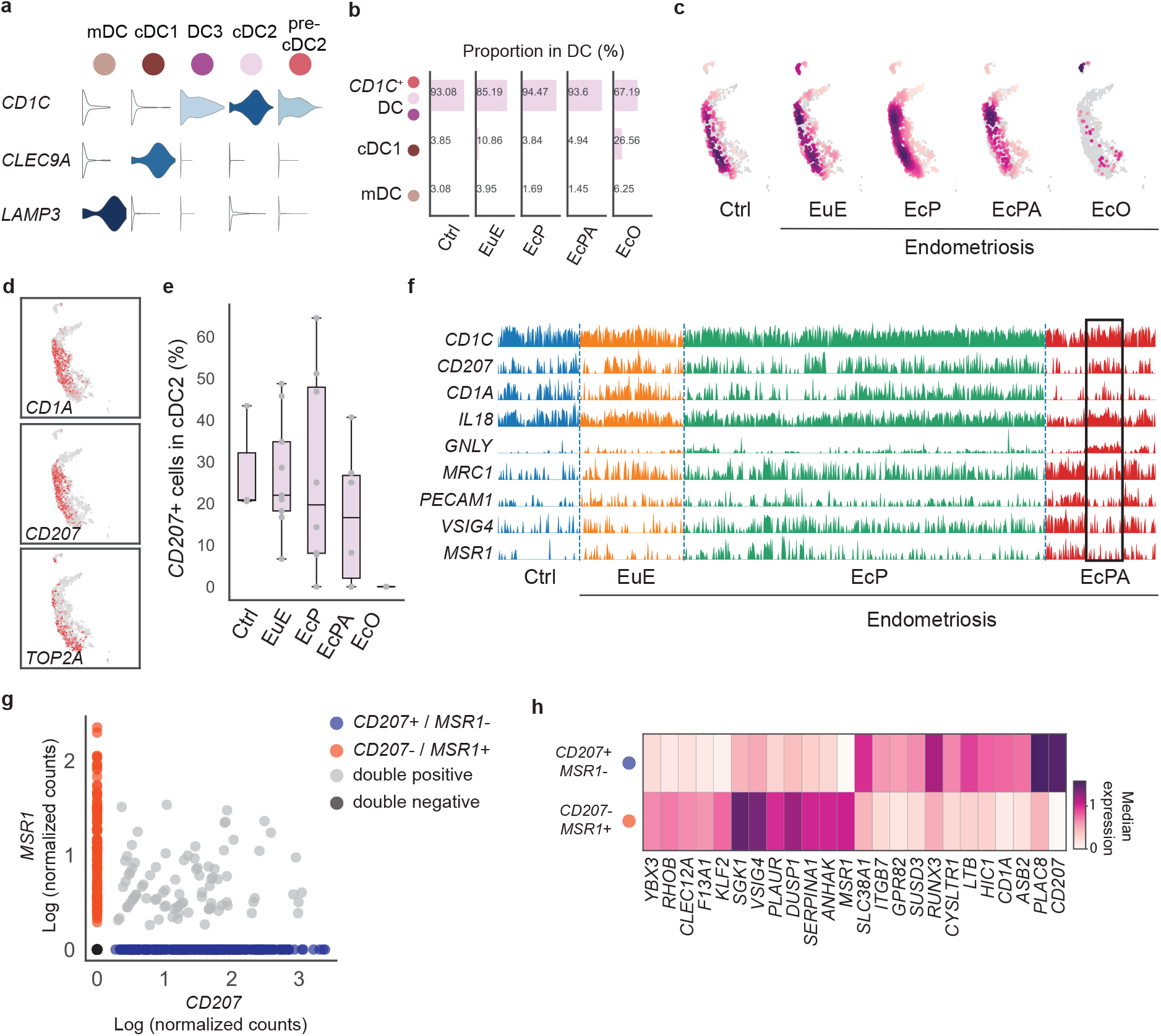
Immunomodulatory role of DC in peritoneal endometriosis. **a**, Violin plot showing markers of dendritic cell (DC) subpopulations, *CD1C* expression is prevalent in three DC subpopulations: pre-cDC2, cDC2 and DC3. **b**, *CD1C^+^*cells represents the majority of the DC population, accounting for more than 85% of DCs in all but ectopic ovary tissue. **c**, Density plot showing the increased cDC2 populations in peritoneal lesion compared to EuE. **d**, Expression of cDC2 markers *CD207* and *CD1A;* and *TOP2A.* **e**, Proportion of *CD207* expressing cells across all cDC2 populations. *CD207*+ cells were consistently observed in eutopic endometrium, but variable in peritoneal lesion and none were observed in ovarian lesions. Each dots represent the percentage of CD207^+^ for each tissue biopsy. The box represents quartiles of the dataset with centerline corresponding to median and whiskers spans the min and max data point **f**, Track plot representing the expression of DEGs upregulated in cDC2-CD1A in EcPA (Wilcoxon, FDR < 0.05). Each bar represents a cell. Differential expression for the represented genes is detected in EcPA cells (black frame). **g**, *S*catter plot showing *CD207+/MSR1-* (n*=* 242), *CD207*-/*MSR1+* (n= 118), double positive (n=88) and double negative (n= 153) cells, and representing different cell states. **h**, Top 12 DEGs between *CD207*+*/MSR1-* and *CD207*-/*MSR1*+ populations in cDC2 subpopulations within EcP and EcPA (Wilcoxon, FDR < 0.05, logFC > 1).

While some studies have also reported altered DC proportions in endometriosis^32, 33^, the field is still lacking a comprehensive characterization of DC heterogeneity across affected tissues. Trajectory analysis suggests that cDC2s derive from pre-DCs, and this transition aligns with the substantial number of proliferative pre-cDC2 cells we observe (Supplementary Fig. 7b,c). The relationship between cDC2s and DC3s appears tissue-specific; in the Ctrl samples, cDC2 and DC3 cells seem to derive from an intermediate population (red arrows, Supplementary Fig. 7b), where DC progenitor- and hematopoietic stem cell-derived blood-derived DC-specific genes (*FLT3, SIGLEC6,* and *AXL*)^34^ are co-expressed (Supplementary Fig. 7d). In EcP, this dynamic is altered and pre-cDC2s appear to be the major source of other cDC2 and DC3 subpopulations. Previous studies have suggested that DCs maintain themselves within a tissue by proliferating under normal conditions, but can be bolstered by an influx of blood-derived DCs during periods of heightened immune activity, such as viral infections^35^.

As a possible reservoir for tissue-resident DC maintenance, we further investigated the diversity among endometriosis-derived cDC2s. A subset of cDC2s in eutopic endometrium and peritoneal lesions, but not ectopic ovary, uniquely express *CD207*, a gene expressed by Langerhans cells and immature DCs that localize in the epidermis and most non-lymphoid tissues^36^ (Fig. 4d-f). Further analysis revealed cellular heterogeneity across cDC2 populations in EcP and EcPA, with two major cell states harboring a dichotomous expression for *CD207* and *MSR1* (Fig. 4f,g). Differential gene analysis highlighted that *CD207*^+^ cDC2 cells express genes related to immunogenic DC maturation (*IL18*, *GNLY*, *RUNX3, LTB*)^37–39^, whereas *MSR1*^+^ cDC2s express immunomodulatory genes (*MRC1*, *VSIG4*, *SGK1,* and *PECAM1*)^40–42^ (Fig. 4h). GSEA analysis of cDC2 across tissues indicated that phagocytosis and cytokine-mediated signaling pathways were upregulated in endometriosis tissues (Supplementary Fig. 8). This demonstrates disease-specific dendritic cell heterogeneity and highlights a potential immunomodulatory role for *MSR1*^+^ cDC2s in the peritoneal microenvironment.

### Analysis of lymphocyte subpopulations reveals endometriosis-specific cell-cell communication axis and organization

Next, we interrogated lymphocyte subpopulation diversity and the interactions of lymphocytes with other immune subpopulations in endometriosis tissue (Fig 5a, Supplementary Fig. 9a). We found numerous interactions between T_Reg_s and various immune subpopulations, particularly with macrophages. One such interaction is between *CCL18* and *CCR8* that is observed specifically in EuE and EcPA. *CCL18*, encodes for a cytokine implicated in chronic inflammatory disease^43^, is differentially expressed in both Mϕ1-LYVE1 and Mϕ5-activated subpopulations (Fig 5b). *CCR8*, encoding a critical driver for T_Reg_-mediated immune tolerance^44^, is exclusively express by T_Reg_ in EuE, EcP and EcPA (Fig. 5b). Through this endometriosis specific interaction, this result suggests that macrophages cooperate with T_reg_s to promote an immunomodulatory microenvironment. Further, we found that genes associated with T_Reg_ regulatory function are altered between control and endometriosis (Fig. 5c): maintenance of the tolerogenic immune function of T_Reg_ in Ctrl is attributed to the expression of *HAVCR2*, *LAG3, ENTPD1, ICOS, TNFRSF4,* and *CTLA4;* while the expression of *TIGIT*, *PRDM1* and *CD96* is prevalent in endometriosis tissues and specifically in EcO. It is also worth noting that *ENTPD1—*which encodes an important regulator in uterine NK cells that promotes immune tolerance and angiogenesis during pregnancy^45^*—*is upregulated in NK1 subpopulation cells in both ectopic lesions (Fig. 5d). Collectively, these changes in gene expression indicate modulation of interactions between the various immune subpopulations in ectopic lesions, though the exact mechanism remains unclear.

**Fig. 5.**
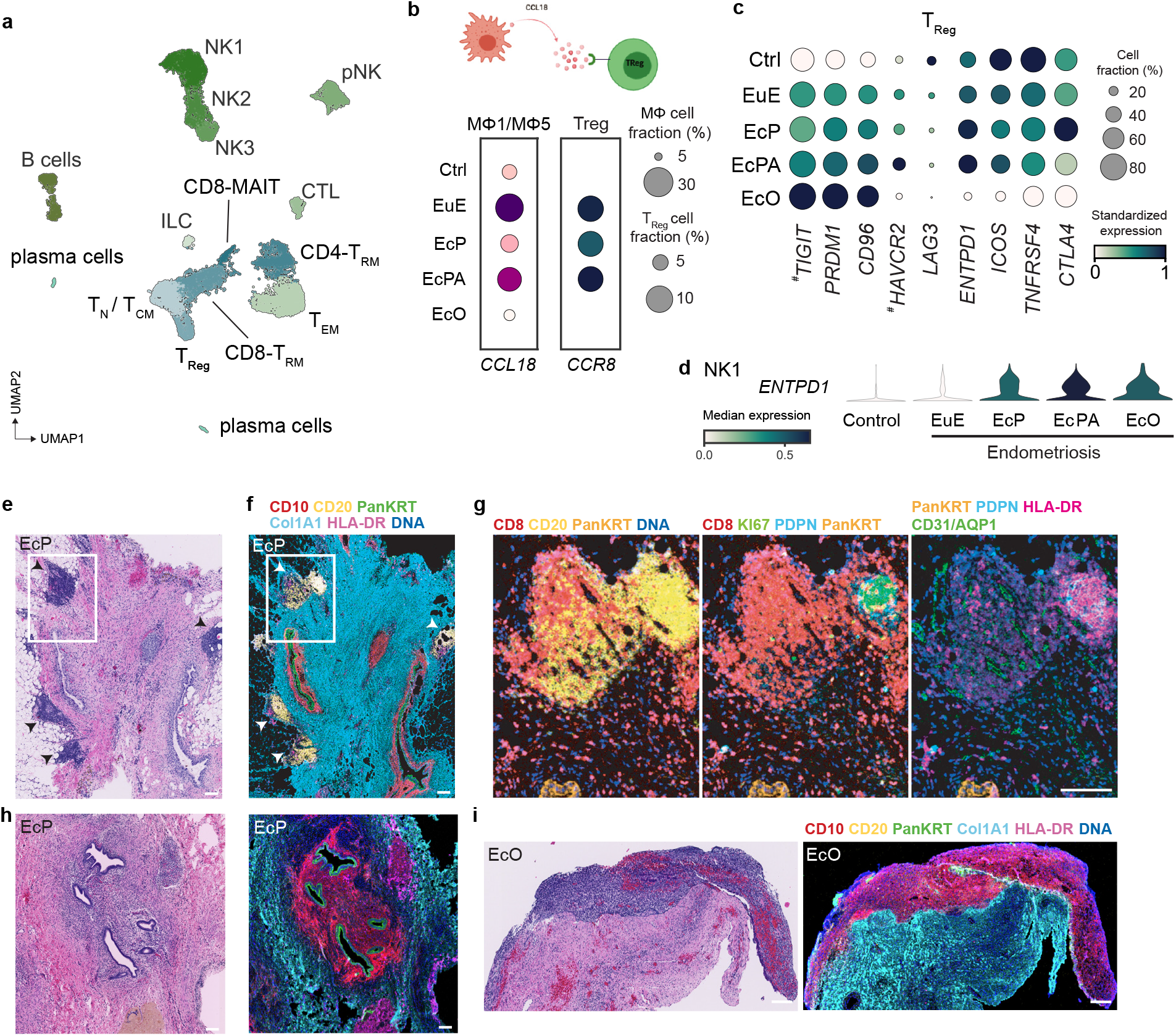
TLS presence in peritoneal endometriosis. **a**, UMAP plot of lymphocyte subpopulations. Represented clustering highlights 14 different subpopulations based on known markers. **b**, Schematic showing CCL18-CCR8 ligand-receptor interaction between macrophages Mϕ5 and Tregs. The dot plot shows gene expression for this interacting pair in each tissue type. **c,** Dot plot showing DEGs associated with TReg self-tolerance maintenance, (^#^ marks non-significant DEG). **d,** Violin plot representing *ENTPD1* gene expression in tissue-resident NK1 cells across sample types. **e**, H/E staining from FFPE tissue section of a peritoneal lesion. This patient sample presented TLS-like formation highlighted in the white frame. **f,** IMC image from the same lesion showing endometrial fibroblasts (CD10), B-cells (CD20), epithelial cells (Pan-KRT), stroma (Col1A1) and antigen presenting cells (HLA-DR). TLS are primarily located through an accumulation of CD20^+^ cells in the periphery of the lesion (white frame). **g**, Magnified image showing GC structures with accumulation of B-cells (CD20) in the center surrounded by T-cells (CD8). KI67 labels proliferative B-cells within the GC. CD31 and AQP1 label blood endothelial cells. PDPN marks follicular dendritic cells. HLA-DR overlap with CD20 indicates an antigen presenting capacity within the GC. **h**-**i**, H/E (left) and corresponding IMC (right) representative images from endometriotic lesions without TLS in EcP (**h**) and EcO (**i**) for identical antibodies panels in (**f**). Scale bar =100 µm.

Using our scRNA-seq-informed IMC panel, we interrogated the immune cell distribution among ectopic lesions. Unexpectedly, we uncovered the presence of immune cell clusters fitting the description of tertiary lymphoid structures (TLS) in peritoneal lesions (Fig. 5e-g). TLSs are characterized by the presence of a germinal center (GC) microarchitecture consisting of a central B-cell cluster surrounded by T-cells among other cell types. TLSs are commonly seen in autoimmune disease, chronic inflammatory disease, and more recently described in various tumor microenvironments but have never been described in endometriosis^46, 47^. We did not detect similar structures in ovarian lesions or across all EcP (Fig. 5h,i), suggesting TLS formation may not be a driver of lesions but perhaps a consequence, in some instances, of a sustained inflammatory response. Gene expression analysis of B cells shows subtle transcriptomic differences among genes related to GC B-cells such as *BCL6, SEMA4A and CXCR5*^47, 48^ (Supplementary Fig. 9b), suggesting that this phenomenon is variable among EcP lesions and between patients.

Altogether, these data emphasize the diversity among immune cells co-existing within endometriosis lesions where the innate immune system actively promotes immune tolerance.

### Characterization of a novel epithelial progenitor-like cell population in eutopic endometrium and ectopic endometriosis

We identified ten epithelial populations composing the epithelial endometrial glands and mesothelium (Fig. 6a,b) some of which show similarity to cell types identified in healthy endometrium^5, 6^ whereas other populations differ substantially or have not been previously observed. In addition to experimental differences, we reasoned that the contraceptive treatment-induced alternate secretory-like state likely accounts for these discrepancies. Of the two previously unreported populations in our dataset, one corresponds to mesothelial cells—found in ectopic tissue—while the other consists of *MUC5B*+ epithelial cells (Fig. 6a). We found *MUC5B*+ population to be present in both eutopic and ectopic tissues and to uniquely express *RUNX3*, *TFF3* and *SAA1* (Fig. 6b, Supplementary Fig. 10a,b). We confirmed the presence of epithelial MUC5B^+^ cells in eutopic endometrium through immunohistochemistry (Fig. 6c). *SAA1* encodes for serum Amyloid A (SAA), a HDL-associated lipoprotein and major modulator of inflammation^49^, epithelial pro-restitution in mucosal wound closure^50^, and promotor of phagocyte chemotaxis through FPR2^51^. This prompted us to look for possible interacting partners for *MUC5B*^+^ cells through ligand-receptor analysis. We found *FPR2*, the gene encoding for the SAA receptor to be uniquely expressed by myeloid cells, particularly monocytes and Mϕ4-infiltrated cells (Supplementary Fig. 10c). Trajectory analysis indicated that *MUC5B*^+^ epithelial cells preceded other endometrial-like epithelial cell populations in both eutopic and ectopic tissues (Fig. 6d), indicating *MUC5B*^+^ epithelial cells as potential progenitor cells.

**Fig. 6.**
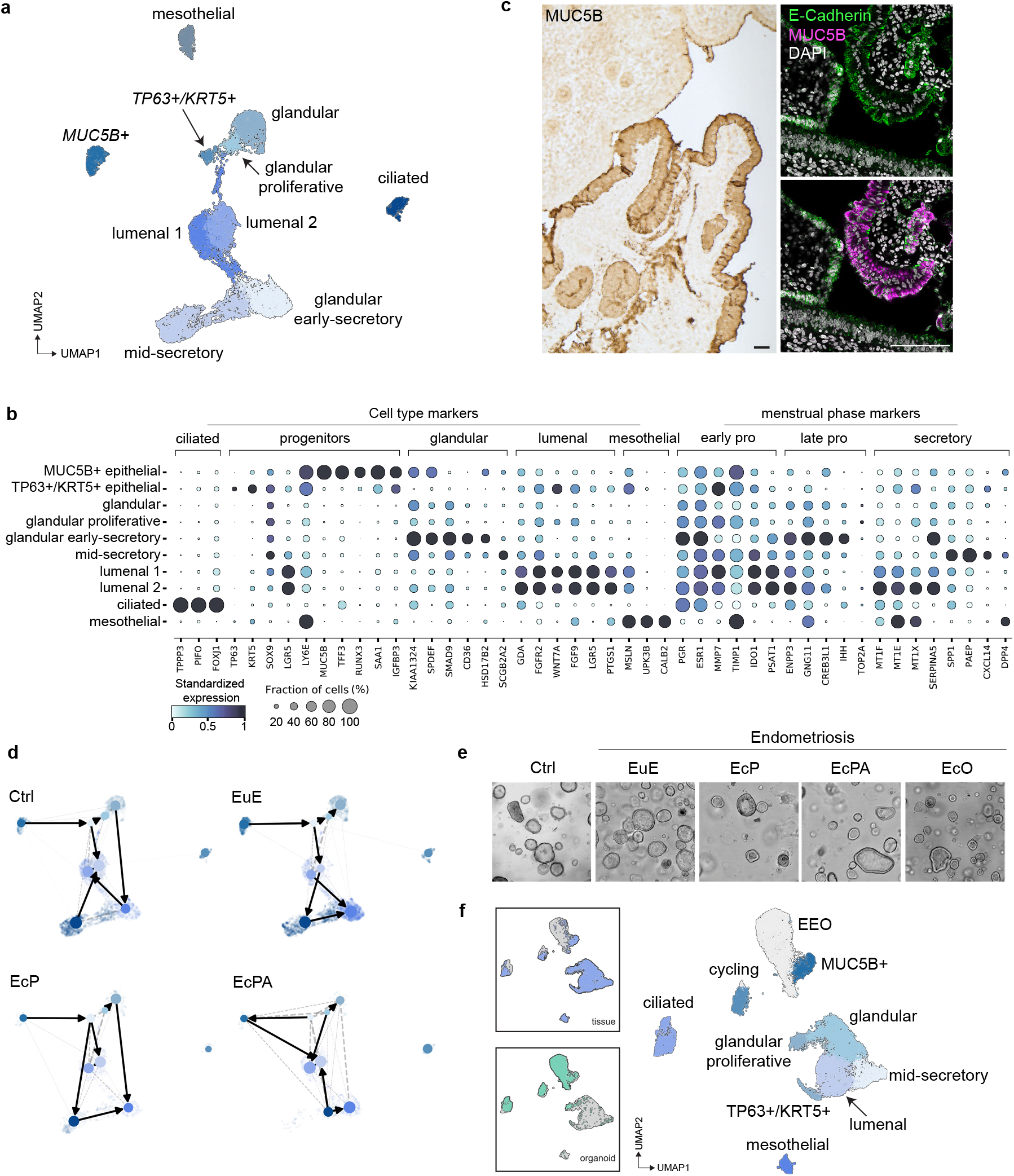
Characterization of epithelial cell subpopulations in Ctrl and endometriosis patients. **a**, Unsupervised clustering of epithelial cells led to 10 subpopulations (n=18,923) represented in the UMAP. **b**, Markers for each epithelial subtype and menstrual phase across each epithelial cell subtype. **c**, Immunohistochemistry (IHC) staining confirms the presence of MUC5B^+^ cells in EuE (Left panel). Immunofluorescence (IF) showing co-localization of endometrial epithelial (E-Cadherin^+^, in green) and MUC5B^+^ cells (magenta). Nuclei were counterstained with DAPI (cyan) in EuE. Scale bar = 100µm. **d**, PAGA trajectory inference from RNA velocity analysis of epithelial cells for each tissue type. Mesothelial cells were excluded from this analysis. We also excluded EcO due to low cell numbers (n=104). **e**, Representative image of endometrial epithelial organoid (EEO) cultures derived from dissociated single-cell of endometrium and endometriotic lesions. **f**, UMAP plot representing the merge dataset for *in vivo* (tissue derived) and *in vitro* (EEO) epithelial cells. We identify 10 clusters through Leiden clustering.

In parallel with scRNA-seq of primary tissue, we derived endometrial epithelial organoids (EEO) from single cells to further interrogate the regenerative capacity of the epithelial glands present in eutopic and ectopic tissues (Fig. 6e). EEOs were maintained in proliferative condition and subsequently profiled using scRNA-seq, yielding a total of 12,757 cells (Supplementary Fig. 11a), which we combined and analyzed together with the single-cell epithelial cell transcriptomes from primary tissue (Fig. 6f). Of the populations identified in the primary tissue, EEO cells mostly distribute along ciliated and proliferative epithelial cells, but also formed a new cluster containing cycling epithelial cells (Fig 6f). The largest population of EEOs cells clusters together with the *MUC5B*^+^ epithelial cells from primary tissue (Fig. 6f) and expresses markers similar to *MUC5B*^+^ cells *in vivo* (Supplementary fig. 11b). Altogether, our data highlighted epithelial cell heterogeneity present in eutopic and ectopic endometria, revealing an uncharacterized endometrial progenitor-like cell population common to all tissues and present in organoid cultures.

## Discussion

We report the first comprehensive description of endometriosis lesions at single-cell resolution and compare this data to single cell transcriptome data we generated from control endometrium, endometrium from endometriosis patients and, organoids derived from these tissues. We analyze both peritoneal and ovarian lesions, the two most common types. Our approach was holistic, to capture the entirety of cell types (or at least those that survive dissociation) that constitute lesions and their adjacent surroundings and thus provide a view of the cellular composition and communication within the niche where lesions establish and evolve. To provide a spatial context to this data we utilize IMC, a hyperplex antibody-based imaging method, with selection of antibodies guided by the scRNA-seq data. Finally, our data is generated from patients under oral contraceptive treatment but with active disease, thus ensuring data interpretation, such as cell-cell communication, is not confounded by normal menstrual cycle dynamics.

Our study details the precise cellular composition of endometriosis tissues. While we identify extensive similarities in cell type composition between the eutopic endometrium to that of peritoneal lesions (i.e. fitting the definition of a lesion being a piece of tissue from the eutopic endometrium), we also detected profound dysregulation of the innate immune and vascular systems in peritoneal lesions. Ovarian lesions, however, display extensive and distinct cell type composition and expression differences to that of peritoneal lesions. Single cell analysis provides important clues on the interconnected cellular networks where myeloid, endothelial, epithelial and pericyte subpopulations influence the formation of the endometriosis-favoring microenvironment.

Among myeloid cell subpopulations, macrophages and DCs have been described as key players in endometriosis pathology^1, 2, 4^ with reports showing endometriosis-related alterations of macrophages^26, 52^ and DCs^32, 33^. However, a comprehensive description of myeloid sub-types that scRNA-seq provides was lacking up until now. We show that macrophages (Mϕ1-LYVE1 and Mϕ4-infiltrated) exhibit an immunotolerant phenotype when compared to the equivalent cell populations in control endometrium. In addition, we describe DCs expressing the mannose receptor *MRC1* and the immunoregulatory *VSIG4* which are capable of promoting immunosurveillance escape and angiogenesis, thus benefiting lesion establishment in endometriosis^53^ and cancer development^54^. We present a precise characterization of immunomodulatory macrophage and DC populations in peritoneal endometriosis that adopt a coordinated immunotolerant phenotype in the endometriosis microenvironment. Such a phenotype was previously reported in decidual macrophages and associated to fetal tolerance during pregnancy^55, 56^. Thus, the present dataset constitutes an ideal starting point to understand how endometriosis may hijack a naturally occurring immunotolerant process to sustain lesion formation and evolution. A deeper understanding of this myeloid compartment (in addition to the lymphoid) in endometriosis is critical, as therapeutics targeting the immune system have been proposed as strategies for treatment^2, 57, 58^. A functional understanding of each myeloid subpopulation’s role will determine if these cells constitute key drivers of the disease, and therefore key therapeutic targets, or simply a byproduct of the continuous inflammation provoked by lesion settlement.

The accumulation of myeloid cells we identify in lesions is interconnected with exacerbated vascularization, a distinctive trait of peritoneal lesions that appears accentuated in the adjacent tissue surrounding the lesions. We report the presence of *CCL19*^+^ (and *CCL21^+^*) perivascular cells. While similar cells were reported in primary and secondary lymphoid organs and shown to play a role in immune cell chemoattraction^59, 60^, this population has not yet been described in endometriosis. In addition to attraction of immune cells, our data suggests these *CCL19*^+^ perivascular cells likely promote angiogenesis through angiopoietin expression, presumably targeting endothelial tip cells in the peritoneal microenvironment that express the corresponding receptors. In further support of these cells playing an active role in lesion vascularization and growth, it has been shown that inhibition of *SNCG* reduces these features in endometriosis^62^, and our data indicate *SNCG* is uniquely expressed by this *CCL19*^+^ perivascular population. The regulation of angiogenesis through secretion of pericyte-derived pro-angiogenic factors was previously demonstrated in *in vitro* proinflammatory microenvironments^63^ and here we illustrate *in vivo* angiogenesis modulation by perivascular cells in peritoneal endometriosis. The peritoneal angiogenic setting contrasts from the ovarian lesion microenvironment where *CCL19*+ perivascular cells are absent. Our data also identifies extensive additional differences between the two types of lesions. Thus, while the endometriosis field encompasses both ovarian and peritoneal lesions under a common disease name and treatment, we provide a holistic understanding of the disease that highlights fundamental lesion type differences that should be taken into account for therapeutic strategy design, such as vascular targeting strategies^64, 65^.

Endometrial epithelial glands form integral components for both eutopic endometrium and endometriotic lesions. The characterization of endometrial epithelial stem cells has been challenging due to the dynamic nature of the regenerative endometrium. Recent single-cell driven descriptions of endometrial epithelial cells from healthy endometrium provide important insights into epithelial subpopulations and the associated hormone responses across the menstrual cycle^5, 6^. The field, however, is still lacking a precise characterization of stem-like epithelial cell populations that could explain epithelial gland establishment and initial lesion formation in ectopic tissues. With the 32,000 epithelial cells we capture in our study, we were able to uncover a novel progenitor-like epithelial cell population expressing *MUC5B* among other specific markers. These cells were detected in all samples, at 2-10% of all epithelial cells within tissues sampled but most abundant (78% of cells) in the organoids, which were maintained in proliferative conditions, a pattern of expression one would expect of a progenitor cell. We confirmed, through gene expression and protein detection, the presence of *MUC5B*+ epithelial cells across patients in both eutopic endometrium and ectopic lesions, suggesting their putative role in ectopic lesions. However, while some marker genes suggest a pro-restitutive role and interactions with myeloid cells, it remains unclear how this cell subset contributes to the cyclic regeneration of endometrium during the menstrual cycle or to ectopic epithelial gland formation, alone or in conjunction with other described progenitor cells found in the endometrium, such as *SOX9*+ cells^6^. While *SOX9* is expressed in our *MUC5B* population it is also broadly expressed across other epithelial sub-types as well. Further functional studies will be key to define their precise role in the endometriosis niche.

Thus, we have generated a comprehensive description of the heterogeneity composing the eutopic endometrium and ectopic ovarian and peritoneal endometriosis lesions. This atlas represents a unique tool to understand the key players and their dynamic interplays that constitute the endometriosis niche. We believe this dataset will be instrumental for designing effective therapeutic strategies or diagnostic biomarkers to provide some relief to the large group of underserved endometriosis patients.

## Methods

### Human endometrium and endometriosis tissue collection

Tissue samples were obtained from the University of Connecticut Health Center (UCHC). All tissue donations and experiments were approved by the Institutional Review Board at UCHC, The Jackson Laboratory, and the Human Research Protection Office of U.S Department of Defense. Pre-menopausal patients (aged 18 to 49 years old) pre-operatively diagnosed with stage II-IV endometriosis and scheduled for laparoscopic surgery were invited to participate in this study. Endometriosis staging was confirmed at the time of laparoscopy according to the revised American Society for Reproductive Medicine guidelines. All patients were treated with oral contraception at the time of sample collection (Supplementary Table 1). Matched eutopic endometrium and endometriosis tissues were collected from endometriosis patients (Fig. 1a; Supplementary Fig. 1a). Eutopic endometrium was obtained by performing an endometrial biopsy during hysteroscopy. Ectopic peritoneal endometriosis was obtained by resecting the entire endometriosis lesion and adjacent peritoneum ensuring the entire visible lesion was excised. For the control cohort, non-endometriosis eutopic endometrium biopsies were obtained from patients scheduled for surgery who were not suspected to have endometriosis. Complete patient demographic information is provided in Supplementary table 1. Upon resection, fresh tissue was immediately stored in MACS tissue storage solution (Miltenyi, 130-100-008) and kept on ice until processing.

### Tissue dissociation for single-cell RNA sequencing

Fresh tissues were immediately processed for scRNA-seq. Ectopic endometriosis lesions from the peritoneum were divided into ectopic lesion (EcP) and ectopic adjacent (EcPA) (Supplementary Fig. 1a). Viable single cells were obtained by mechanical and enzymatic digestion using cold active protease (CAP), following a modified version of the previously described protocol (Adam 2017). Briefly, minced tissue was transferred to GentleMACS C tubes (Miltenyi, 130-096-334) containing protease solution (10mg/ml *Bacillus Licheniformis* protease (CAP) (Sigma, P5380) in DPBS supplemented with 5mM CaCl2 and 125U/ml DNaseI (Stemcell, 07900) and incubated in cold water bath (6°C) for 7-10 minutes, performing trituration steps every 2 minutes. After incubation, sample were mechanically dissociated on a Miltenyi GentleMACS Dissociator for 1 minute, twice. Undigested tissue was allowed to settle by gravity for one minute. Single cells within the supernatant were transferred into a collection tube containing wash buffer PBS supplemented with 10% fetal bovine serum (FBS) (Gibco, 10082147), 2mM EDTA, and 2% bovine serum albumin (BSA, Miltenyi 130-091-376). Remaining undissociated tissue was incubated with fresh CAP protease for a total of 20 to 40 minutes, proceeding with a trituration step every 5 minutes and a Miltenyi gentleMACS Dissociator step every 15 minutes. After recovery of single cells, residual undissociated tissue was incubated with PBS supplemented with 1 mg/ml dispase on the Miltenyi gentleMACS Dissociator at 37°C for 15 minutes, and until complete tissue dissociation. Single cells were then pelleted, washed, and filtered through 70µm MACS Smartstrainer (Miltenyi, 130-098-462). Prior to FACS sorting, single cell suspension were stained with propidium iodide (PI) (BD Biosciences, 556364) and calcein violet (Invitrogen, C34858) in FACS buffer (PBS, 2mM EDTA, 2% BSA) and according to manufacturer protocols. Viable cells (propidium iodide negative and calcein violet positive) were sorted using the BD FACS Aria Fusion cell sorter and recovered in Advanced DMEM/F12 (Gibco, 12634010) supplemented with 2mM GlutaMAX (Gibco, 35050061), 10mM HEPES (Gibco, 15630080), 20% FBS, 1% BSA. Sorted viable cells were then washed and resuspended with 0.04% BSA in PBS and assessed for viability using trypan blue staining for subsequent scRNA-seq experiments.

### Endometrial epithelial organoid cultures and cell-hashing for scRNA-seq

Following tissue dissociation and single cell recovery, and after 10x chromium chip loading, remaining single cells were pelleted and resuspended in cold Matrigel (Corning, 356231). Fifty microliter (50 µL) droplet were plated onto 24-well plate wells (Greiner Bio-one, 662102) to generate endometrial epithelial organoids (EEOs). After Matrigel dome solidification, organoid media was added to cover each dome, as previously describe by Boretto *et al.*^66^. Organoid passaging was performed every 7-10 days and according to the established protocol from Turco *et al.*^67^. For scRNAseq experiments, organoid cultures between passage 3 and 5 and at day 7-11 after plating were collected, washed twice with wash media (Advanced DMEM/F12, 2mM GlutaMAX, 10mM HEPES, 0.1% BSA), and dissociated into single cells using TrypLE Express (Gibco, 12605010) for 3-5 minutes at 37°C. Cell suspensions were filtered with 40 µm mesh filter to remove debris and cell aggregates. Lastly, cells were washed and resuspended with cell staining buffer (Biolegend, 420201) for hashing with TotalSeq-A anti-human Hashtag reagents (Supplementary Table 6, Biolegend) for 30 min at 4°C and following previously published protocol^68^. After staining, cells were washed to remove excess antibody and resuspended in PBS/0.04% BSA for subsequent counting. Hashed cells were assessed for viability and sorted for viable cells as described below.

### Single-cell capture, library preparation, and sequencing

Single cell suspensions were analyzed for viability and counted on a Countess II automated cell counter (Thermo Fisher). A total of 12,000 cells were loaded onto a channel of 10X Chromium microfluidic chips for a targeted cell recovery of 6,000 cells per lane. Single cell capture, barcoding, and library preparation were performed using 10X Chromium v3 chemistry according to manufacturer’s protocol (10x Genomics, CG000183). Sample cDNA and library quality controls were performed using the Agilent 4200 TapeStation instrument and quantified by qPCR (Kapa Biosystems/Roche). Libraries were sequenced on a NovaSeq 6000 (Illumina) with the S2 100 cycle kit targeting 100,000 reads per cell for tissues or 50,000 reads per cell for organoids.

### Single-cell data preprocessing and clustering

Illumina base call files for all libraries were demultiplexed and converted to FASTQs using bcl2fastq v2.20.0.422 (Illumina). The CellRanger pipeline (10x Genomics, version 3.1.0) was used to align reads to the human reference GRCh38.p13 (GRCh38 10x Genomics reference 3.0.0), deduplicate reads, call cells, and generate cell by gene digital counts matrices for each library. The resultant counts matrices were further processed with Scanpy package (version 1.7.1)^69^ to exclude genes that are detected in less than 3 cells and to exclude cells with (1) fewer than 500 genes, (2) fewer than 1,000 UMIs, (3) maximum of 100,000 UMIs, and (4) maximum mitochondrial content of 25%. Doublet identification were performed using Scrublet^70^. Filtered matrices were then combined and normalized such that the number of UMI in each cell is equal to the median UMI across the dataset and log transformed. Scanpy was used to identify the top 2,000 highly variable genes from log transformed combined matrix. The (1) mitochondrial genes, (2) hemoglobin genes, (3) ribosomal genes, (4) cell cycle genes^71^, and (5) stress response genes were excluded from highly variable gene set^72^. Principal component analysis and neighborhood graph generation were performed based on highly variable genes set. Harmony (version 1.0) batch correction was performed to reduce variabilities introduced by inherent patient differences, tissue types, and endometriosis staging to enhance clustering by cell type^73^. Batched-corrected principal components were used for dimensionality reduction using Uniform Manifold Approximation and Projection (UMAP). Clustering was then performed with Leiden community detection algorithm^74, 75^. Further doublet identification was calculated based on the median distance of a cell to the center of its respective cluster centroid in UMAP space and the coexpression of marker genes of two or more cell types. All suspected doublets were removed from the analysis.

### Cell types and cell state identification

Marker genes of each cluster were identified using Wilcoxon Rank-Sum test in a one-versus-rest fashion, with (1) minimum 0.5 - 2 fold change between group, (2) expressed by at least 0.7 fraction of cells in the group, and (3) expressed by maximum 0.3 fraction of cells outside the group. Cell types were determined by matching the biomarkers with previously described cell types and cell states, and from biomarkers curated from the literature.

### Comparative analysis with Bulk RNA-seq

Total RNA of endometrium and endometriotic lesions was extracted from snap frozen tissue or RNAlater (Invitrogen, #AM7020) stabilized tissue using QIAGEN RNeasy Mini Kit according to the manufacturer’s instructions (Supplementary Table 1). Library preparation was performed using KAPA mRNA Hyperprep kit (Roche) according to manufacturer’s instruction. Bulk RNAseq libraries were sequenced on NovaSeq 6000 (Illumina) with SP 100 cycles single-end reads kit resulting in an average of 42.7 million reads per sample. Reads were aligned to the GRCh38.p13 reference genome (GRCh38 10x Genomics reference 3.0.0), filtered, and quantified with nf-core/rnaseq (version 1.4.2)^76^ utilizing the STAR aligner. Read counts were normalized to counts per millions (CPM) reads. scRNA-seq was compared to bulk RNAseq by utilizing pseudo-bulk transform (summing UMI counts for all cells in each sample and CPM normalization. Differential gene expression between scRNA-seq and bulk RNA-seq data was analyzed with edgeR exactTest^77^. Differentially expressed genes (DEGs) were generated sequentially for eutopic endometrium (Control and EuE), ectopic peritoneal endometriosis (EcP and EcPA), and ectopic ovarian endometriosis (EcO).

### Identification of DEGs and GSEA analysis between tissue typ**es**

DEG analysis between tissue types within a population was performed on clusters with more than 500 cells. We utilized edgeR’s glmQLFTest function to compare each tissue types to Control samples. Significant DEGs were considered at FDR < 0.01 (Supplementary Table 4). Gene Set Enrichment Analysis (GSEA) to GO Biological process (2018) was performed on significant DEG (FDR < 0.00001) with gseapy prerank function for each cell subtype. Resulting enriched gene ontology list was filtered at FDR < 0.10 (Supplementary Table 5).

### Correlation matrix, dendrogram, cell cycle phase and cell density estimation

Analyses were executed with functions implemented in Scanpy (1.7.1) package. Similarities between eutopic endometrium (Control and EuE) tissues were based on hierarchical clustering calculated from Pearson correlation using the Ward linkage algorithm. Cell cycle phase (G1, S or G2M) estimation was calculated following the protocol previously described in Satija *et al*.^78^ and based on markers retrieved from Tirosh *et al*.^79^. The cell density was estimated with Gaussian kernel density estimation on major cell subtype within each tissue type.

### Trajectory Inference

Read counts of spliced and unspliced RNA was computed with velocyto (0.17.17)^80^ on all 10x libraries obtained from tissue biopsies. We utilized the *run10x* function which takes output from CellRanger pipeline. Reads were aligned to GRCh38 (10x Genomics reference 3.0.0) and GRCh38 repeat mask downloaded from UCSC Genome Browser as recommended. Projected stream and PAGA trajectory was calculated with scVelo (0.2.3) following recommended workflow previously described^81, 82^. First, clusters of interest are isolated based on cell barcodes (e.g., myeloid cells in Control). Second, spliced and unspliced counts were log normalized and used for nearest-neighbors estimation. Then, RNA velocity was computed using scVelo’s dynamical model which infers the splicing trajectory for each gene and allows for differential kinetics across distinct lineages and functional states that may be present in the dataset. Lastly, the velocity was visualized via streamlines and PAGA graph abstraction to visualize the general trajectory of each cell cluster.

### Ligand-receptor analysis

Ligand-receptor analysis was performed using CellPhoneDB (2.1.4)^83^ on all 58 subclusters. We modified the protocol by running CellPhoneDB on each 10x library separately to reflect the interactions only within individual tissue sample. As such, we added additional parameters to obtain list of interactions that are (1) p-value < 0.01, (2) detected in at least 50% fraction of each tissue type, (3) is not self-interaction, and (4) is a unique cell-to-cell interactions (number of cell type pair is less than 150 counts).

### Histology and immunofluorescence

Formalin-fixed paraffin-embedded (FFPE) tissues were cut into 5-μm sections, mounted on slides and stained for hematoxylin and eosin (H/E). The slides were then scanned with a Hamamatsu Nanozoomer slide scanner for histopathological examination. Immunofluorescence staining was performed on FFPE tissue sections. Slides were incubated for 10 minutes at 55°C in a dry oven, deparaffinized in fresh Histoclear (National Diagnostics, #HS-200), and rehydrated through a series of graded alcohols. Antigen retrieval was performed in a decloaking chamber (BioSB TintoRetriever) for 15 minutes at 95°C in neutral citrate buffer, pH 6.00 (Abcam, #ab93678). Tissue was blocked and permeabilized with 10% donkey serum/0.1% Triton X-100 in PBS for 30 minutes at room temperature, then incubated with primary antibodies MUC5B (1/1000, Novus Biologicals, #NBP1-92151) and E-cadherin (5 µg/ml, R&D Systems, #AF648) overnight. Tissue sections were subsequently incubated with secondary antibody Donkey anti-rabbit Alexa Fluor 647 (Invitrogen, #A-31573) and Donkey anti-Goat Alexa Fluor 488 (Invitrogen, #A-11055) for 1 hour at room temperature. DAPI (1 µg/ml, Sigma, MBD0015) was used to counterstain the nuclei, then mounted with ProLong Diamond (Thermo Fisher, #P36970). Images were taken using a Leica SP8 Confocal microscope at 40x magnification and processed with FIJI^84^.

### Image Mass Cytometry (IMC)

FFPE of EcP and EcO tissues were cut into 5-μm sections and mounted on slides. Slides were incubated for 15 minutes at 55°C in a dry oven, deparaffinized in fresh histoclear, and rehydrated through a series of graded alcohols. Antigen retrieval was performed in a decloaking chamber (BioSB TintoRetriever) for 15 minutes at 95°C in citrate buffer, pH 6.0. After blocking in buffer containing 3% BSA, slides were incubated overnight at 4°C with a cocktail of metal-conjugated IMC-validated primary antibodies and described in Supplementary table 6. The following day, slides were washed twice in DPBS and counterstained with iridium intercalator (0.25 μmol/L) for 5 minutes at room temperature to visualize the DNA. After a final wash in ddH20, the slides were air-dried for 20 minutes. The slides were then loaded on the Fluidigm Hyperion imaging mass cytometer. Regions of interest were selected using the acquisition software and ablated by the Hyperion. The resulting images were exported as 16-bit “.tiff” files using the Fluidigm MCDViewer software and analyzed using the open source Histocat++ toolbox/ Histocat web^85^.

### Statistics and reproducibility

All hypothesis tests were conducted with the Wilcoxon rank-sum test unless otherwise stated, and the Benjamini-Hochberg correction was used to correct for multiple simultaneous hypotheses tests where applicable.

## Acknowledgments

We thank the following Jackson Laboratory (JAX) Scientific Services cores, partially supported through the JAX Cancer Center Support Grant (CCSG) P30CA034196-30, for expert technical assistance: Single Cell Biology, Flow Cytometry and A. Carcio and T. Prosio, Genome Technologies and R. Maurya, Histology, and Microscopy. We also thank the JAX cyberinfrastructure team for computational resources, L. Perpetua and the UConn Health Research Biorepository, and the UConn Health Surgery Center Personnel for assistance in biopsies collection. We would like to thank the Clinical and Translational Research Support group, the Sponsored Research Administration and the Research Program Development services for administrative assistance. All schematic panels were created with Biorender.com. This study was supported by the Department of Defense Congressionally Directed Medical Research Programs (CDMRP) Discovery Award Grant W81XWH1910130 (E.T.C), JAX Institutional startup funds (P.R.).

## Author Contributions

E.T.C., P.R. and D.E.L. conceived and designed the study. Y.T. and E.T.C., performed and supervised the experiments. Y.T. and E.T.C. optimized the tissue dissociation protocol. Y.T., D.L. and S.B.B. performed tissue dissociation and single-cell experiments. D.E.L. and A.A.L. consented patients and collected clinical samples. S.S. performed IMC experiments. Y.T., W.F.F., and C.E.T. performed data analysis. Y.T. ,F.W.F. and E.T.C. wrote the manuscript. Y.T., W.F.F., A.A.L., P.R., D.E.L. and E.T.C., contributed critical data interpretation. All authors have read or provided comments on the manuscript.

