## Supplementary files for "Single cell analysis of endometriosis reveals a coordinated transcriptional program driving immunotolerance and angiogenesis across eutopic and ectopic tissues"

Supplementary Figure 1

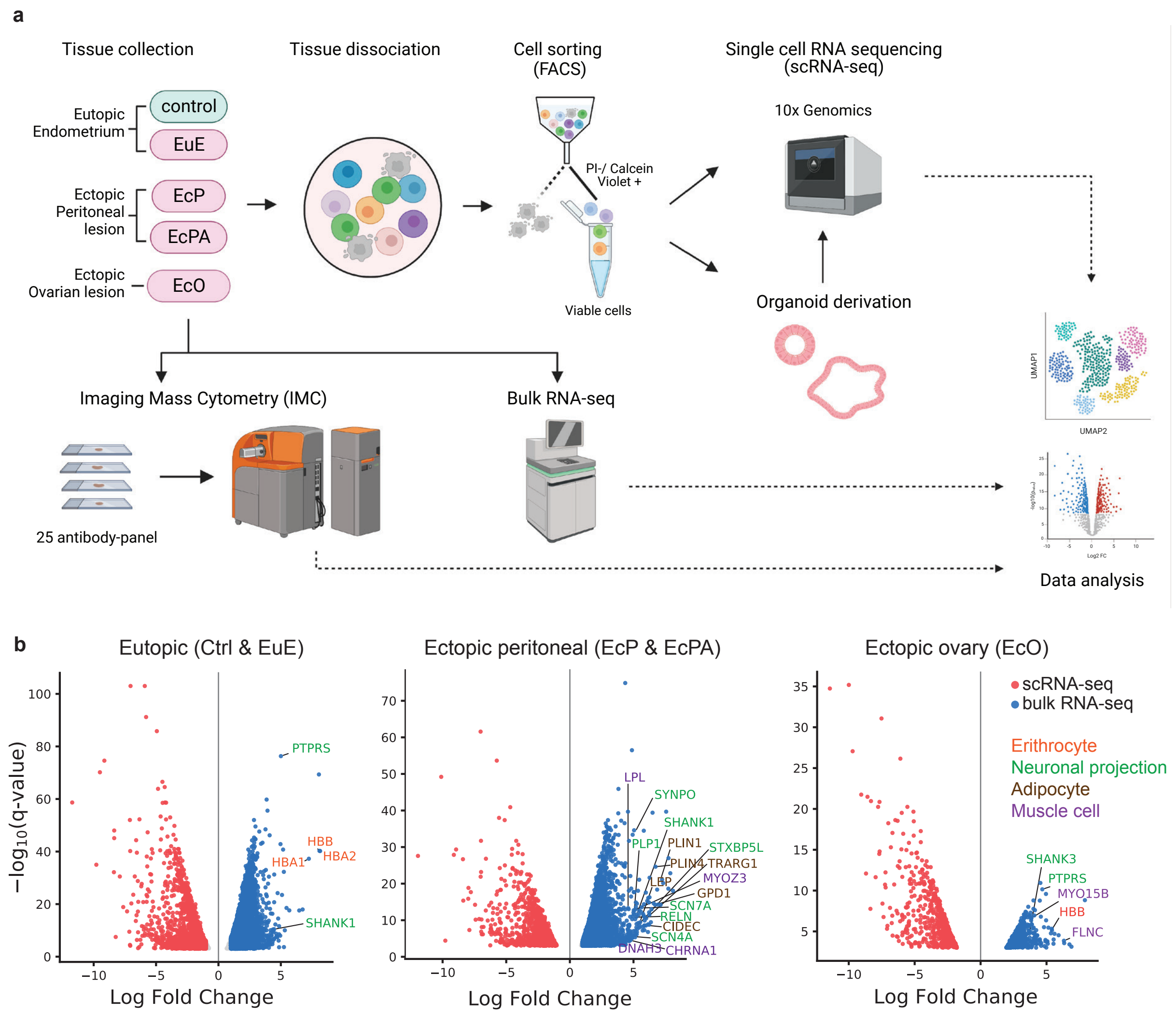

**Supplementary Fig. 1. | Overview of experiment design and bulk RNA-seq profiles from Ctrl and endometriosis patient. a**, Experimental workflow. **b**, Volcano plots representing DEGs between scRNA-seq pseudo bulk (red) and bulk RNA-seq from undissociated tissue (blue). The genes highlighted are exclusively expressed in bulk RNA-seq and associated with erythrocytes (orange), neuronal projections (green), adipocytes (brown), and muscle cells (purple).

Supplementary Figure 2

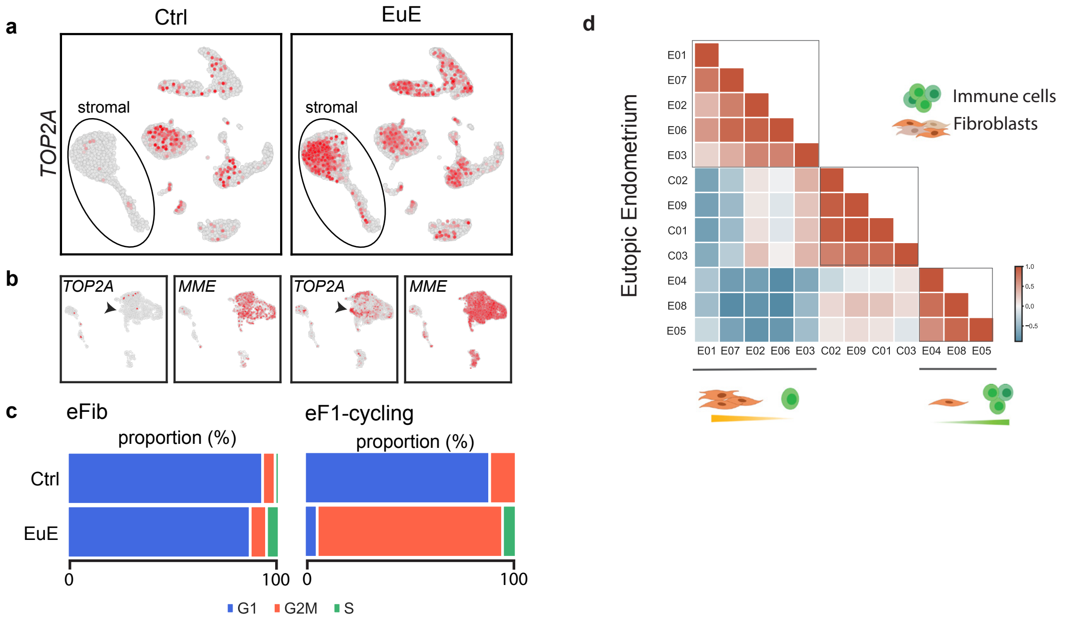

**Supplementary Fig. 2. | Cellular composition of Ctrl and endometriosis eutopic endometrium. a,** UMAP representation for the expression of *TOP2A* in Ctrl and EuE in the main clustering. Circle denotes the stromal cell population. **b,** Representation of *TOP2A* (marker for proliferating cells) and *MME* (marking endometrial fibroblasts) expressing cells in Ctrl and EuE stromal cell subclusters. Arrows depict a subpopulation expressing *TOP2A* in EuE, subsequently named as eF1-cycling. **c,** Proportion of cells in G1, G2M, S cell cycle phases within eFib and eF1-cycling subpopulations. **d**, Matrix plot representing the overall similarity of endometrium biopsies from control and endometriosis (Pearson correlation based on gene expression from each patient). EuE clustered into two groups, each one showing an enrichment for fibroblast or immune cells.

Supplementary Figure 3

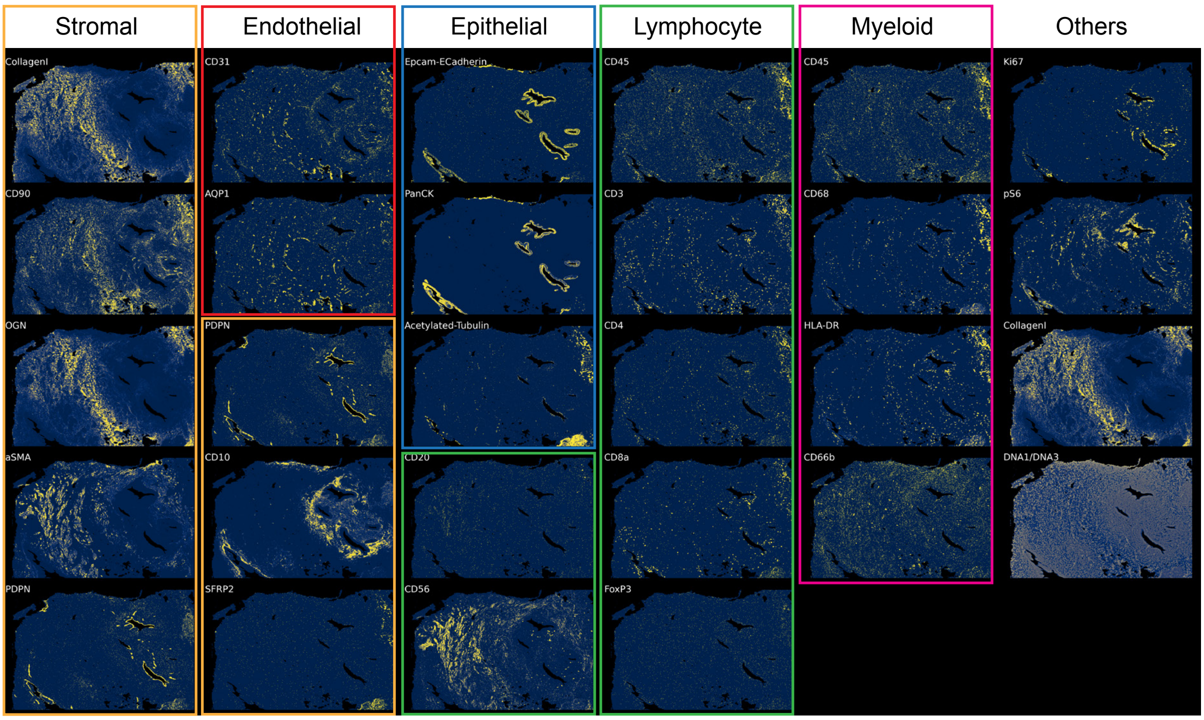

**Supplementary Fig. 3. | IMC panel for spatial profiling of ectopic lesions.** Each antibody was selected according to the cell types inferred by the scRNA-seq data analysis. Images show single channels for each metal-conjugated antibody in a peritoneal endometriosis lesion. A total of 23 antibodies was used to identify cellular heterogeneity within stromal, endothelial, epithelial, lymphocyte, and myeloid major cell types. Additional antibodies (in "Others") were used to identify cell proliferation (Ki67), active metabolism (pS6), extracellular matrix (Collagen1), and nuclei (DNA1/DNA3).

Supplementary Figure 4

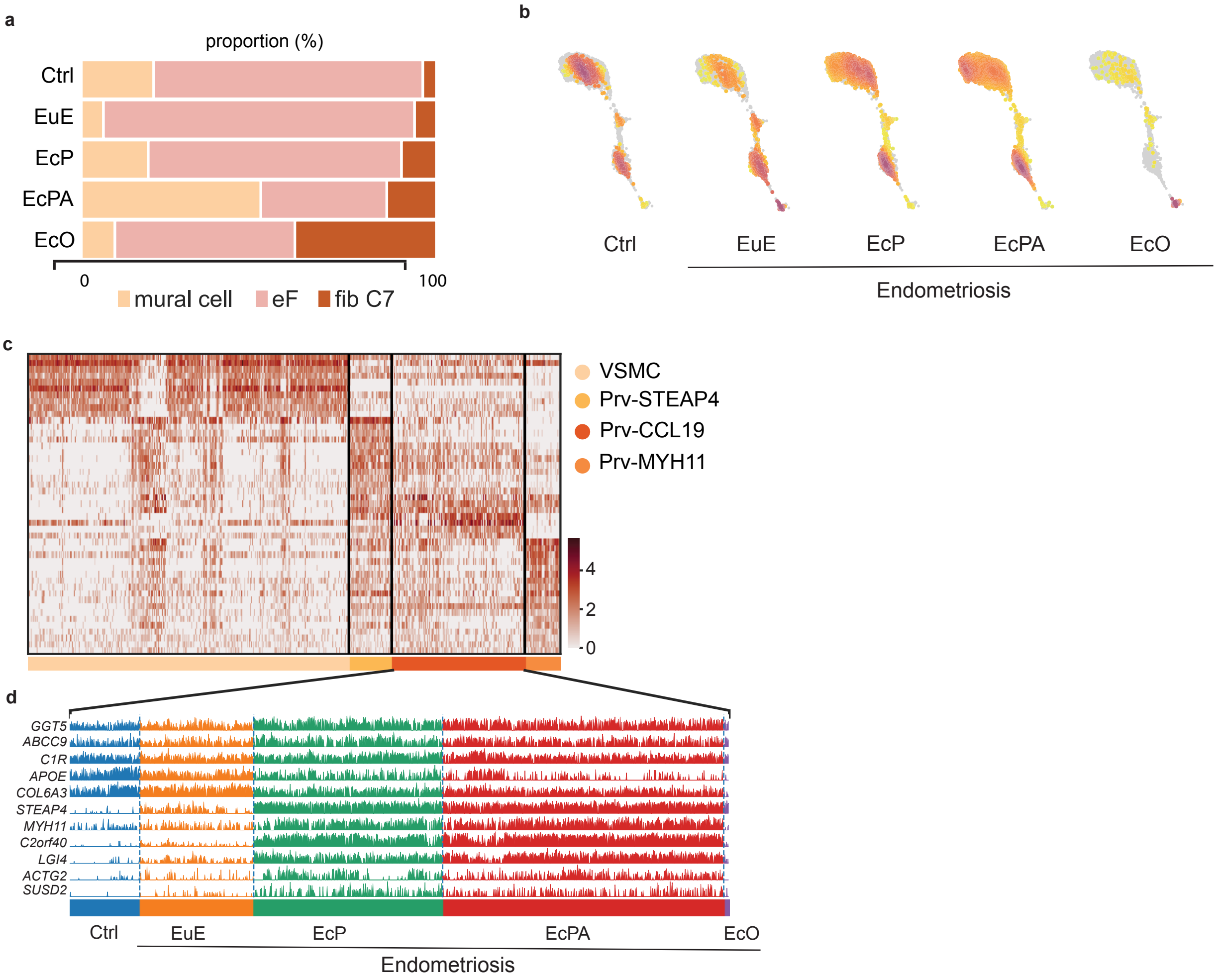

**Supplementary Fig 4. | Stromal cell analysis across sample types. a**, bar plot representing proportion of stromal cell types in control endometrium and endometriosis lesions. Endometrial fibroblasts were found in all lesions. Fibroblast C7 is the predominant fibroblast type in EcO. **b**, Density plot showing distribution of mural cells for each tissue. **c**, Heatmap of markers genes for mural cell subtypes. **d**, Track plot representing gene expression pattern for selected DEG in Prv-CCL19 subpopulations. *GGT5* and *ABCC9* are pan-markers for this cell subtype.

Supplementary Figure 5

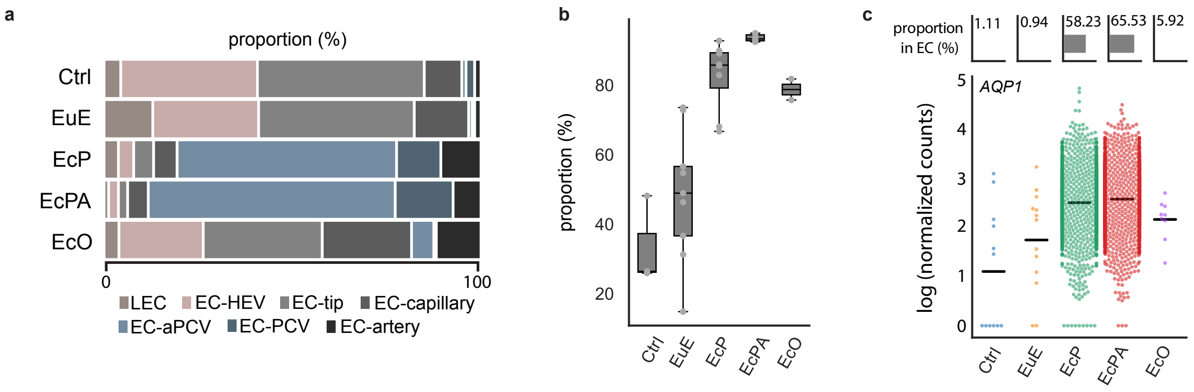

**Supplementary Fig. 5. | Characterization of endothelial cells (EC) across sample types**. **a**, EC proportions by sample type. **b**, EC-aPCV and EC-PCV cell abundances are substantially increased in peritoneal lesions (EcP and EcPA). **c**, (top) Proportion of aPCV among ECs across tissue types. (bottom) Swarm plot showing *AQP1* expression per cell. Horizontal lines represent the median value.

Supplementary Fig. 6

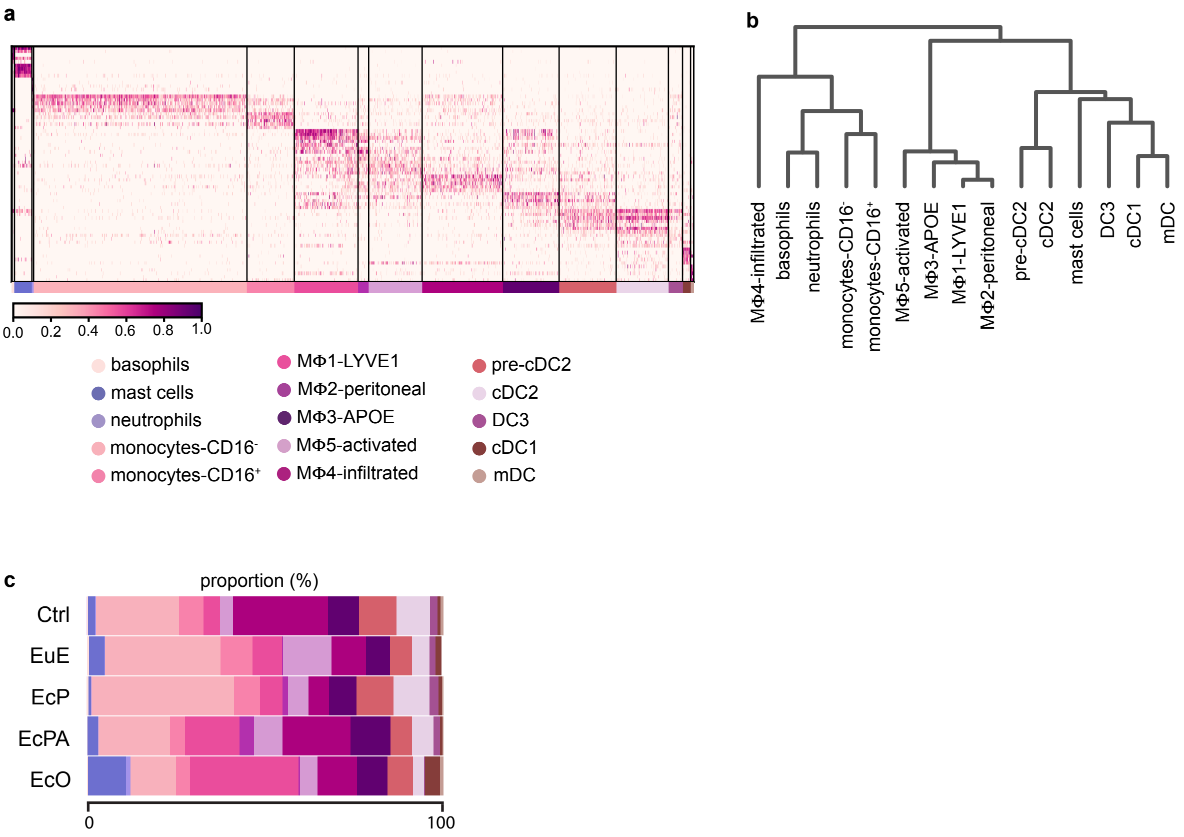

**Supplementary Fig. 6. | Myeloid cell diversity in control and endometriosis. a, Heatmap representing** marker genes for each myeloid subpopulation. **b,** Dendrogram showing the hierarchical clustering (Pearson correlation) for the myeloid cell clusters. **c,** Bar plot showing the representation of each myeloid subtype across tissue types.

Supplementary Figure 7

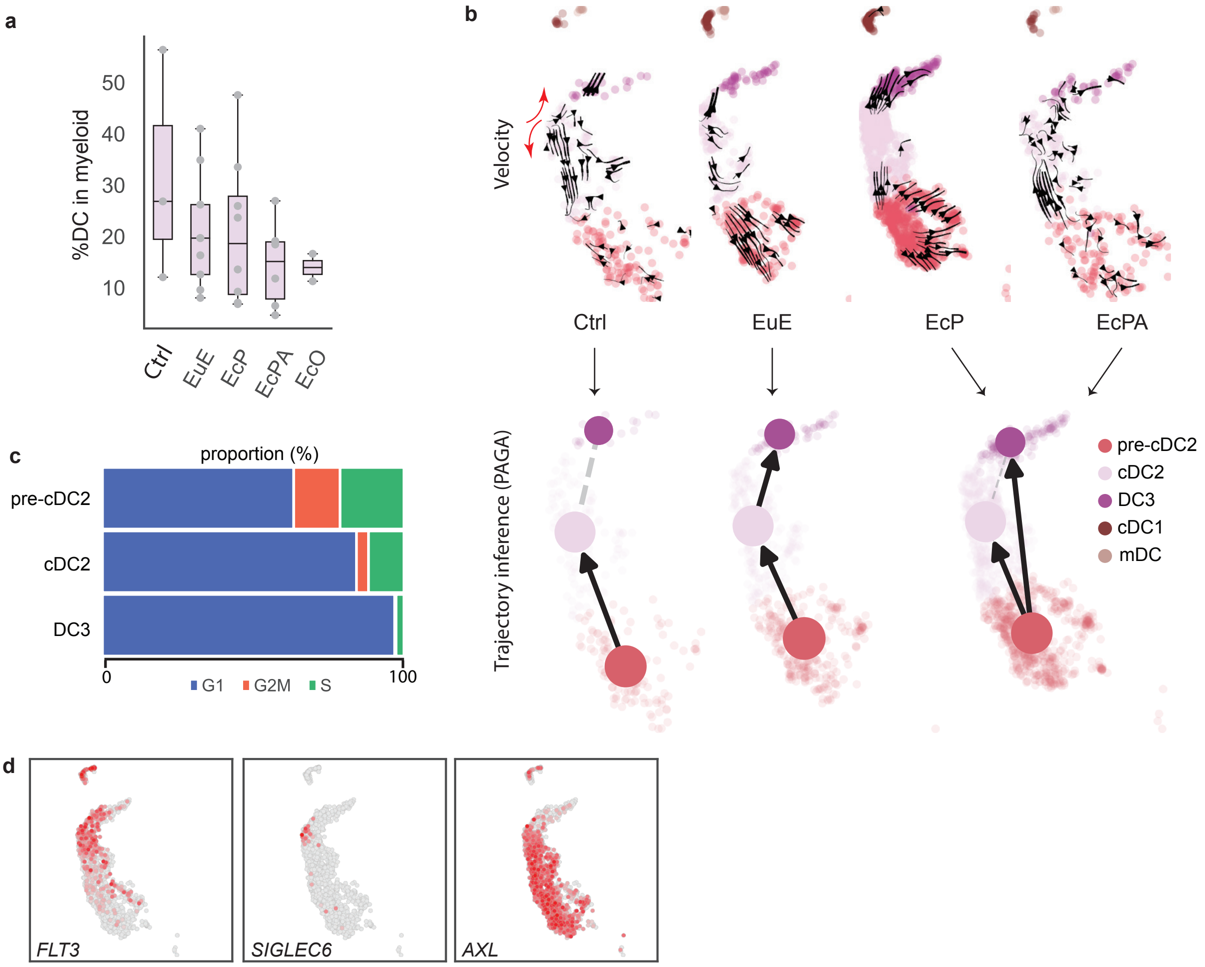

**Supplementary Fig. 7 | DC subpopulations. a**, Bar plot represents the proportion of DCs among all myeloid cells for each patient. Patient-to-patient variability was observed in DC proportions within the myeloid population and across different tissue types. **b**, PAGA and RNA velocity trajectory analysis suggests that pre-cDC2 differentiate towards cDC2 and DC3 in Ctrl and EuE. Red arrows indicate that some cDC2 and DC3 cells derive from a smaller intermediate cell population. **c**, Cell cycle analysis for pre-cDC2, cDC2 and DC3 populations. **d**, Expression of DC progenitor markers *FLT3*, *AXL*, and *SIGLEC6*.

Supplementary Figure 8

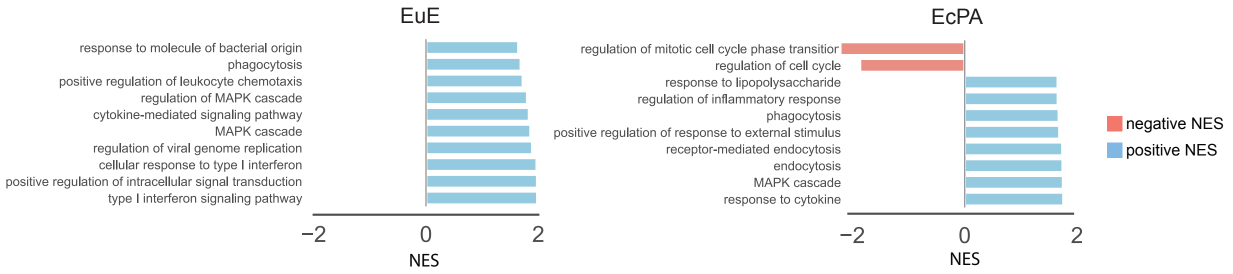

**Supplementary Fig. 8. | Phagocytosis pathway is enriched in cDC2 subpopulations of peritoneal lesions.** Bar plot shows the Normalized Enrichment Score (NES) for the top 10-GSEA pathways in cDC2 cells of EuE and EcPA (FDR < 0.1).

### Supplementary Figure 9

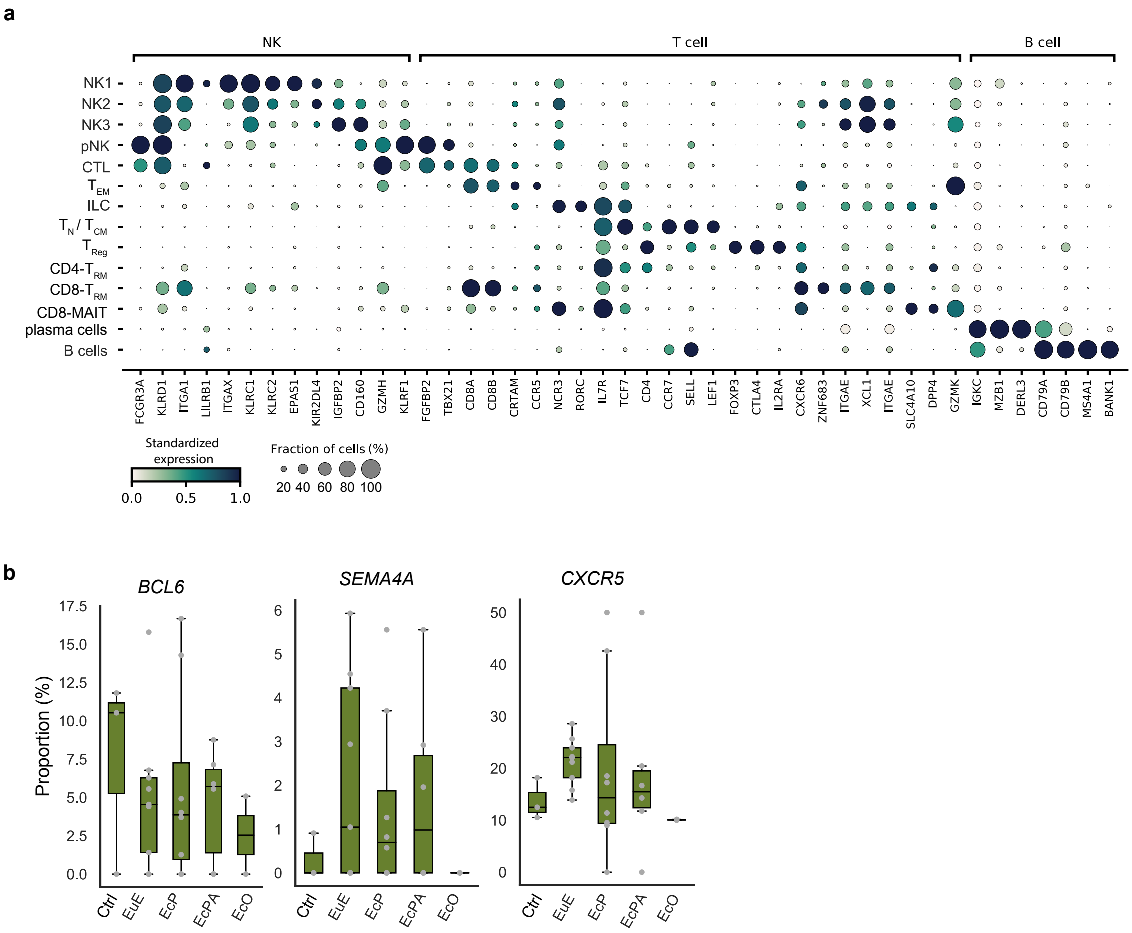

**Supplementary Fig. 9. | Lymphocyte subpopulations in control and endometriosis tissues. a,** Dot plot representing marker genes for each lymphocyte subpopulation, including four natural killer cell (NK) clusters, innate lymphoid cells (ILCs), effector memory T-cells ($T_{EM}$), cytotoxic T-lymphocytes (CTL), naïve/central memory T-cells ($T_N$/$T_{CM}$), T regulatory cells ($T_{Reg}$), CD4- and CD8- tissue resident T cell (CD4-$T_{RM}$ and CD8-$T_{RM}$, respectively), CD8 mucosal-associated invariant T cells (CD8-MAIT), plasma cells, and B cells. **b,** Proportion bar plot of *BCL6*, *SEMA4A*, *CXCR5* expressing cells from the total B cells within each sample type. Each dot represents a unique patient and bar plots represent median distributions.

Supplementary Figure 10

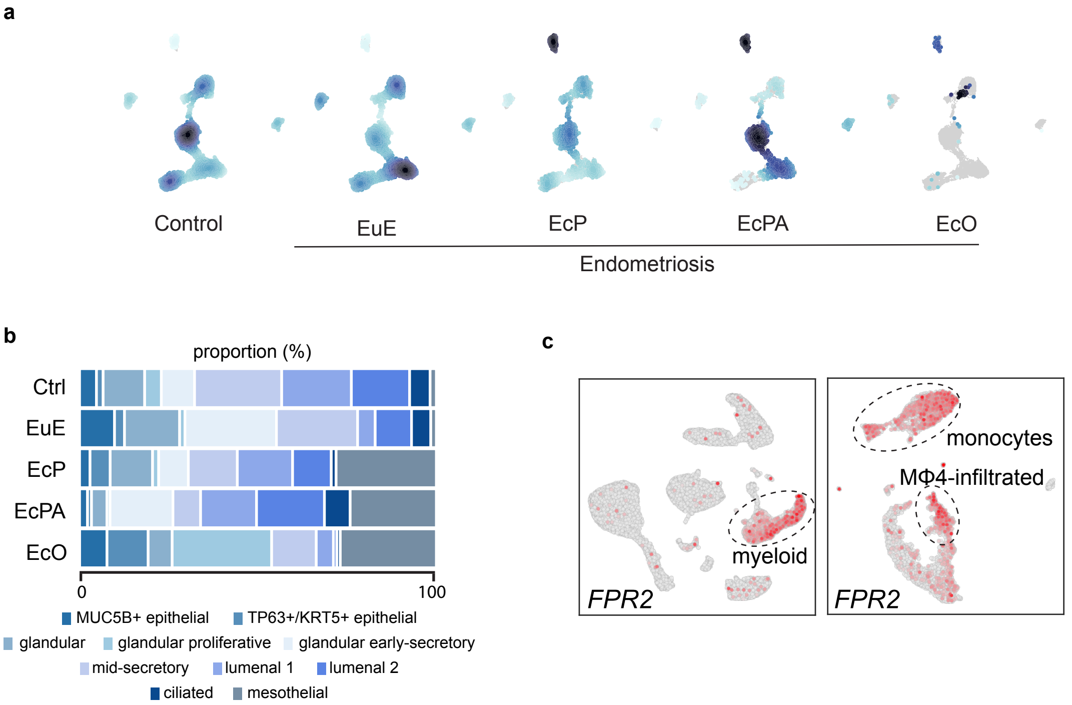

**Supplementary Fig. 10 | Characterization of *in vivo* epithelial cells. a**, Density plot showing the distribution of epithelial subtypes across tissues. **b**, Proportions of epithelial cell subpopulations per sample type. Mesothelial cells appear in EcP, EcPA, and EcO. **c,** Formyl Peptide Receptor 2 (*FPR2)* expression is specific to myeloid cells (left), and more precisely to monocytes and Mϕ4-infiltrated cells (right).

Supplementary Figure 11

a

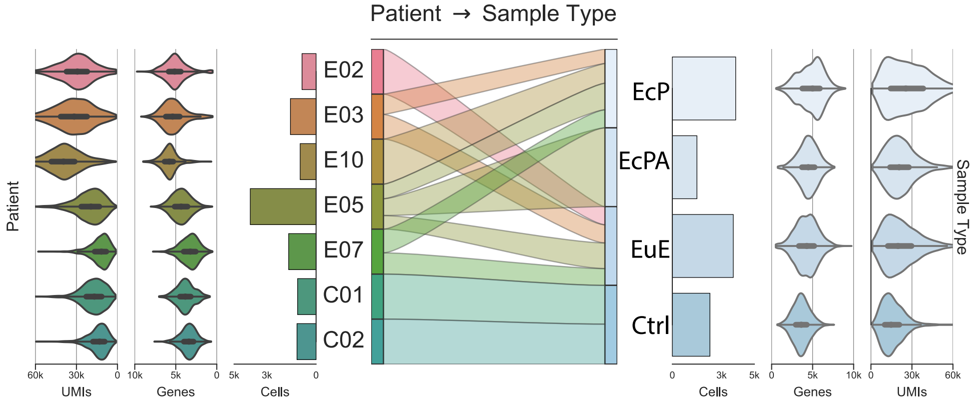


b

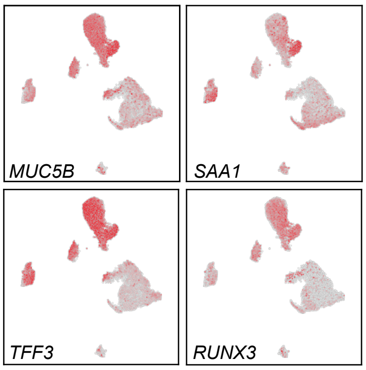

**Supplementary Fig. 11 | Endometrial epithelial organoid (EEO) cells. a**, Sequencing metrics from EEO scRNA-seq representing UMIs and unique genes counts for Control (C) and endometriosis (E) patients and across tissue type. **b**, UMAP showing the co-expression of *MUC5B*, *SAA1*, *TFF3*, and *RUNX3* in the *MUC5B*+ cell population comprising *in vivo* epithelial cells and EEO.

**Supplementary table 1. Patient demographic and experiment performed**

| PID | Age | Ethnicity | Pregnancy | Cohort | rASRM Stage | Drug Treatment | Tissue scRNA-seq | Organoid derivation | Organoid scRNA-seq | IMC | bulk RNA-seq |
|---|---|---|---|---|---|---|---|---|---|---|---|
| E01 | 40 | white | 0 | Endometriosis | 3 | liraglutide, lisinipril, januria | EuE, EcP, EcPA | EuE | | | |
| E02 | 40 | hispanic | 0 | Endometriosis | 2 | norethindrone/E2:1/20 | EuE, EcP, EcPA | EuE, EcP | EuE | | |
| E03 | 33 | hispanic | 0 | Endometriosis | 4 | dostinex, clonazepam, lamictal, seroquel, sumatriptan, norethindrone/E2:1/20 | EuE, EcP, EcPA | EuE, EcP | EuE,EcP | | EuE |
| E04 | 37 | white | 0 | Endometriosis | 4 | mirena, lexapro, trazadone,lorazepam,norethindrone/E2:1/20 | EuE, EcP, EcPA | EuE, EcP | | | |
| E05 | 33 | white | 0 | Endometriosis | 4 | norethindrone/E2:1/20 | EuE, EcP, EcPA | EuE, EcP, EcPA | EuE,EcP,EcPA | EcP | EcP |
| E06 | 30 | white | 0 | Endometriosis | 4 | norethindrone/E2:1/10 | EuE, EcP | EuE, EcP | | | |
| E07 | 34 | white | 0 | Endometriosis | 4 | norethindrone/E2:1/10 | EuE, EcP, EcO | EuE, EcP, EcO | EuE,EcP | EcO | EuE, EcO |
| E08 | 40 | white | 0 | Endometriosis | 4 | norethindrone/e2:1/20 | EuE | EuE, EcP, EcO | | | |
| E09 | 45 | hispanic | 2 | Endometriosis | 4 | norethindrone/e2:1/20 | EuE, EcP, EcPA, EcO | EuE, EcP, EcPA, EcO | | | EuE |
| C01 | 26 | asian | 1 | Control | 0 | norethindrone/E2:1/20, levothyroxine | Control | Control | Control | | Control |
| C02 | 22 | white | 0 | Control | 0 | norethindrone/E2:1/20, lexapro | Control | Control | Control | | |
| C03 | 42 | white | 0 | Control | 0 | n.d. | Control | Control | | | |
| E10 | 34 | white | 0 | Endometriosis | 4 | norethindrone/E2:1/20 | | EuE, EcP | EcP | | |
| E11 | 39 | white | 0 | Endometriosis | 4 | norethindrone/E2:1/20, levothyroxine, lamictal, wellbutrin | | | | EcP, EcO | EuE, EcP, EcO |
| E12 | 33 | black | 0 | Endometriosis | 4 | norethindrone/E2:1/20 | | | | EcP, EcPA, EcO | EcP,EcO |
| C04 | 34 | hispanic | 0 | control | 0 | none | | | | | Control |

Note:

n.d. not disclosed

**Supplementary Table 2. Markers.xlsx (separate file)**

Excel file containing markers of. Each subpopulation defined in this study.

**Supplementary Table 3. DEG-scRNA-seq vs bulk RNA-seq.xlsx (separate file)**

Excel file containing differentially expressed genes between scRNA-seq dataset and bulk RNA-seq dataset in three biopsies type; eutopic endometrium, ectopic peritoneal, and ectopic ovarian.

**Supplementary Table 4. DEG between biopsy type within each subpopulation (separate file)**

Excel file containing differentially expressed genes between endometriosis biopsies type ( EuE, EcP, EcPA, and EcO) and control endometrium (Ctrl), in each of 44 subpopulations defined in this study.

**Supplementary Table 5. GSEA.xlsx (separate file)**

Excel file containing enriched gene ontology term in endometriosis biopsies type (EuE, EcP, EcPA, and EcO) compared to control endometrium (Ctrl), in each of 44 subpopulations defined in this study.

**Supplementary Table 6. Additional information about reagents and antibody used in this study**

Hastags antibodies

| PID | Sample Name | sc-GE Library / sc-hastag Library | Human Hashtag Oligos | Catalogue Number |
|---|---|---|---|---|
| E02 | E02-EuE-o | EOR01 / EOR02 | TotalSeq-A0251 anti-human Hashtag 1 Antibody | 394601 |
| E03 | E03-EuE-o1 | EOR01 / EOR02 | TotalSeq-A0253 anti-human Hashtag 2 Antibody | 394603 |
| E03 | E03-EuE-o2 | EOR01 / EOR02 | TotalSeq-A0253 anti-human Hashtag 5 Antibody | 394609 |
| E03 | E03-EcP-o1 | EOR01 / EOR02 | TotalSeq-A0253 anti-human Hashtag 3 Antibody | 394605 |
| E03 | E03-EcP-o2 | EOR01 / EOR02 | TotalSeq-A0253 anti-human Hashtag 6 Antibody | 394611 |
| E10 | E10-EcP-o | EOR01 / EOR02 | TotalSeq-A0254 anti-human Hashtag 4 Antibody | 394607 |
| E05 | E05-EuE-o | EOR03 / EOR04 | TotalSeq-A0251 anti-human Hashtag 1 Antibody | 394601 |
| E05 | E05-EcP-o | EOR03 / EOR04 | TotalSeq-A0252 anti-human Hashtag 2 Antibody | 394603 |
| E05 | E05-EcPA-o | EOR03 / EOR04 | TotalSeq-A0253 anti-human Hashtag 3 Antibody | 394605 |
| E07 | E07-EuE-o | EOR03 / EOR04 | TotalSeq-A0254 anti-human Hashtag 4 Antibody | 394607 |
| E07 | E07-EcP-o | EOR03 / EOR04 | TotalSeq-A0255 anti-human Hashtag 5 Antibody | 394609 |
| C01 | C01-EuC-o | EOR03 / EOR04 | TotalSeq-A0256 anti-human Hashtag 6 Antibody | 394611 |
| C02 | C02-EuC-o | EOR03 / EOR04 | TotalSeq-A0258 anti-human Hashtag 8 Antibody | 394615 |

IMC antibodies

| Target | Metal | Clone | Supplier | Catalouge Number | Custom Conjugation |
|---|---|---|---|---|---|
| aSMA | 141Pr | 1A4 | Fluidigm | 3141017D | |
| CD10 | 145Nd | Poly | RnD Systems | AF1182 | X |
| CD31 | 146Nd | JC/70A | Novus Bio | NB600-562 | X |
| CD56 | 148Nd | MRQ-42 | Cell Marque | 156R-94 | X |
| SFRP2 | 149Sm | 80.8.6 | EMD Millipore | MABC539 | X |
| CD20 | 151Eu | H1 | BD Biosciences | 555677 | X |
| CD90 | 152Sm | 2D7D11 | Proteintech | 66766-1-Ig | X |
| Ac-tubulin | 154Sm | 7E5H8 | Proteintech | 66200-1-Ig | X |
| CD4 | 156Gd | EPR6855 | Fluidigm | 3156033D | |
| E-Cadherin | 158Gd | 4A2 | Novus Bio | NBP2-54587 | X |
| Epcam | 158Gd | 9C4 | Biolegend | 324229 | X |
| HLA-DR | 159Tb | LN3 | Biolegend | 327002 | X |
| CD8a | 160Gd | C8/144B | Biolegend | 372902 | X |
| CD68 | 162Dy | KP1 | Thermo Fisher Scientific | 14-0688-82 | X |
| CD45 | 163Dy | CD45-2B11 | Thermo Fisher Scientific | 14-9457-82 | X |
| OGN | 164Dy | 2A5F11 | Thermo Fisher Scientific | 66382-1-IG | X |
| FoxP3 | 165Ho | 236A/E7 | Abcam | ab96048 | X |
| PDPN | 167Er | D2-40 | Biolegend | 916606 | X |
| Ki67 | 168Er | B56 | BD Biosciences | 556003 | X |
| Collagen I | 169Tm | Poly | EMD Millipore | AB758 | X |
| CD3 | 170Er | 3F3A1 | Proteintech | 60181-1-Ig | X |
| pS6 | 172Yb | N7548 | Fluidigm | 3172008A | |
| AQP1 | 173Yb | Poly | Thermo Fisher Scientific | PA5-80346 | X |
| Pan Keratin | 174Yb | AE-1/AE-3 | Biolegend | 914204 | X |
